# Curation, inference, and assessment of a globally reconstructed gene regulatory network for *Streptomyces coelicolor*

**DOI:** 10.1101/2021.11.26.470080

**Authors:** Andrea Zorro-Aranda, Juan Miguel Escorcia-Rodríguez, José Kenyi González-Kise, Julio Augusto Freyre-González

## Abstract

*Streptomyces coelicolor* A3(2) is a model microorganism for the study of Streptomycetes, antibiotic production, and secondary metabolism in general. Even though *S. coelicolor* has an outstanding variety of regulators among bacteria, little effort to globally study its transcription has been made. We manually curated 29 years of literature and databases to assemble a meta-curated experimentally-validated gene regulatory network (GRN) with 5386 genes and 9707 regulatory interactions (∼41% of the total expected interactions). This provides the most extensive and up-to-date reconstruction available for the regulatory circuitry of this organism. Only ∼6% (534/9707) are supported by experiments confirming the binding of the transcription factor to the upstream region of the target gene, the so-called “strong” evidence. While for the remaining interactions there is no confirmation of direct binding. To tackle network incompleteness, we performed network inference using several methods (including two proposed here) for motif identification in DNA sequences and GRN inference from transcriptomics. Further, we contrasted the structural properties and functional architecture of the networks to assess the reliability of the predictions, finding the inference from DNA sequence data to be the most trustworthy approach. Finally, we show two applications of the inferred and the curated networks. The inference allowed us to propose novel transcription factors for the key *Streptomyces* antibiotic regulatory proteins (SARPs). The curated network allowed us to study the conservation of the system-level components between *S. coelicolor and Corynebacterium glutamicum*. There we identified the basal machinery as the common signature between the two organisms. The curated networks were deposited in Abasy Atlas (https://abasy.ccg.unam.mx/) while the inferences are available as Supplementary Material.

## Background

Streptomycetes, the largest genus within the actinomycetes, are biotechnologically relevant organisms. They produce around half of the natural antibiotics in current use^1^. However, according to the analysis of genome mining, less than 10% of antibiotics that could be produced by actinomycetes are currently used^2^. Their production could be enhanced not only by experimental technologies such as genetic manipulation but also by a deeper knowledge of their secondary metabolism and transcriptional regulation. *Streptomyces coelicolor* A3(2) has become the model microorganism for the study of antibiotic production and secondary metabolism in general^3^. Before its sequencing, it was already known that *S. coelicolor* produces the red-pigmented antibiotic undecylprodigiosin (RED), the blue-pigmented actinorhodin (ACT), and the calcium-dependent antibiotic (CDA). However, its sequencing revealed more than 20 biosynthetic gene clusters (BGCs). Most of the metabolites produced by these clusters and their regulation are still unknown^4^.

*S. coelicolor* secondary metabolism regulation is very complex. It is controlled by a network of regulators at many levels, from global to cluster situated regulators (CSRs). Most CSRs control their own BGC, however, some of them can bind to multiple BGCs causing a cross-cluster regulation^4^. Sequencing of *S. coelicolor* A3(2) revealed 7825 genes, 965 of them (∼12.33%) code for proteins with a predicted regulatory function. From those, 65 genes coding for sigma factors, a remarkably high number among bacteria, of which ∼70% (45/65) are ECF (extra-cytoplasmic function) sigma factors, suggesting independent regulation of diverse stress response regulons^5^. Besides, it counts with many two-component systems (TCSs), 85 sensor kinases, and 79 response regulators, also related to stress response^5^. The difference between sensor kinases and response regulators suggests a cross-talking among them. Noteworthy, *S. coelicolor* genome codes for several putative regulators that do not belong to families outside *S. coelicolor*^5^. Because of the complexity of the secondary metabolism regulation, a proper understanding of the *S. coelicolor* regulation requires it to be studied systematically at both local and global scales. On a global scale, GRNs are used to study transcription regulation. They can be represented as a directed graph where nodes represent genes, and edges represent the regulatory interactions among the transcription factors (TFs) and their target genes (TGs). Previous comprehensive reviews have been focused on specific morphological differentiation and metabolic processes^4,6-10^. However, a GRN at a global scale is still missing.

The initial approach to reconstruct a global-scale GRN will be through text mining^11^. There we would be able to collect all the information available on the literature for the microorganism. Nevertheless, it would still require manual intervention for those articles where interactions are not clearly defined. Moreover, all genes have not been studied experimentally. Therefore, alternatively, GRN inference has been applied in diverse bacteria to provide a deeper understanding of their regulatory mechanisms. Besides, it has also been applied to propose selective experimental validation of putative interactions, analyze bacterial GRN evolution, and build biological models for biotechnological processes^12-16^. A GRN inference for *S. coelicolor* was performed by Castro-Melchor, et. al in 2010 using ARACNE and applying module validation through the identification of consensus DNA sequences^17^. However, the resulting network was not assessed with any gold standard (GS) available at the time, and no thorough study of its structural properties was performed. Moreover, benchmarking studies of network inference methods have shown the poor predictive power of using a single GRN inference tool^18^.

Here, we performed a collection and curation of the experimentally-validated transcriptional regulatory interactions of *S. coelicolor* A3(2) and classified them based on the confidence level of their supporting evidence. Further, we integrated this curated GRN with previous curations from DBSCR (http://dbscr.hgc.jp/) and Abasy Atlas^19^. Then, we applied the natural decomposition approach (NDA) to identify their system-level components and unveiled different biology aspects of *S. coelicolor* regulation. Next, we applied several tools to infer novel interactions, three based on DNA binding sites for the TFs, and five based on gene expression along with two modifications proposed by the authors. We integrated the predictions using a community approach, which has been reported as the best strategy to reduce the number of false positives^18^. Then we used the most reliable curated network as a GS for the validation of the inferred GRNs. We further assessed the inferred networks through their structural properties and found that the NDA ^20^ is a valuable tool for GRNs dissection and comparison. From the best-rated inferred network, we proposed new TF candidates for the direct regulation of some of the key *Streptomyces* antibiotic regulatory proteins (SARPs) in *S. coelicolor*. Finally, we applied the meta-curated network of *S. coelicolor* to study the conservation of the system-level components with those of its phylogenetically related *C. glutamicum* as an application of the curated network. The workflow of this work and the suggested use of the data herein reported are summarized in Figure 1.

**Figure 1.**
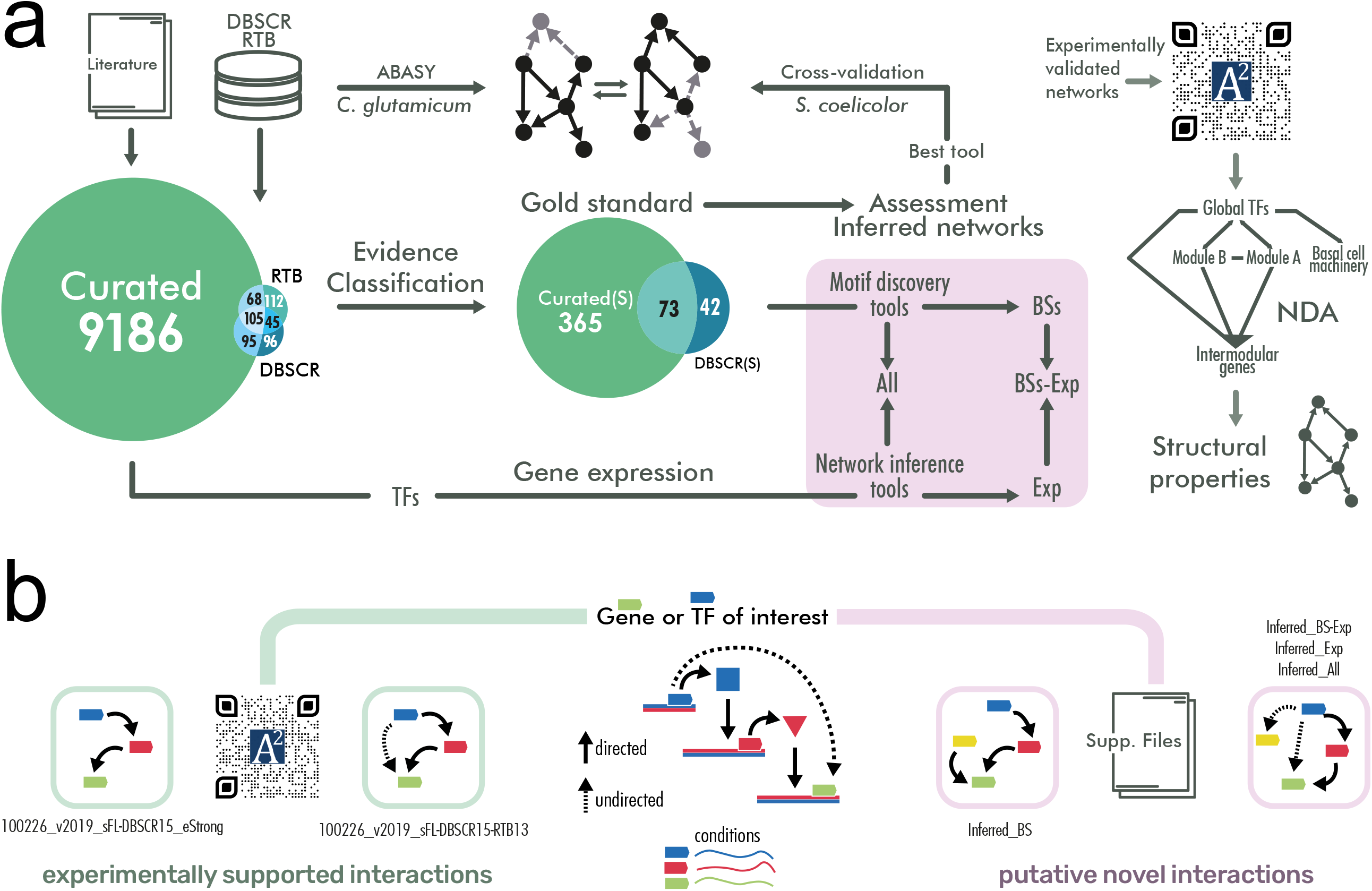
a) Workflow of this work. The purple area covers the inference of the networks. b) Type of interactions contained in the networks. The green path connects to curated regulations supported by experimental evidence. The *“strong”* network contains only the interactions that are supported by an experiment proving that the transcription factor binds a DNA site near a target gene to regulate its transcription. Curated networks without the *“strong”* label might contain indirect interactions, as they could be supported by non-directed experiments (such as gene knockout and its effect on genes transcription). The purple path connects to inferred interactions. Predictions based solely on binding sites predictions would be inferring only TF-DNA interactions. Predictions involving gene expression data might contain indirect interactions.

## Results and discussion

### Reconstruction of the most complete experimentally-validated regulatory network for *S. coelicolor*

We curated a total of 124 papers retrieved from PubMed and Google Scholar queries covering a span of 29 years (from 1990 to July of 2019) (see Figure 2 and Supplementary Table 1). We collected a total of 9714 regulatory interactions (out of the 23908 expected interactions in the complete GRN as predicted by Abasy Atlas v2.4) among 5331 genes. We perceive a notable increment in the number of papers and interactions after the *S. coelicolor* genome was completely sequenced (2002)^5^. This eases the study of its genome and regulation, being 2012 the year with most publications (see Figure 2). We classified the interactions according to their experimental evidence, expanding the RegulonDB scheme^21,22^. First, we label the interactions as “strong” or as “weak” according to the methodology of the experiment performed. A “strong” evidence level is assigned to experiments that prove a physical regulatory interaction between the TF and the TG. This means that the TF can bind to the upstream region of the regulated gene. Here we have experiments such as EMSA in purified proteins or *in vitro* transcription assays. On the other hand, a “weak” evidence level is assigned when there is no evidence of direct interaction. This means that the experiment suggests either a hypothetical DNA binding site, such as ChIP; or an effect of the TF over the gene that might be indirect, through another TF, such as microarray, RNA-Seq, RT-PCR, etc. For experiments that were not in the RegulonDB scheme, such as DNA-affinity capture assay (DACA)^23^, we analyzed their methodology to classify them either as “strong” or “weak” evidence. Supplementary File 2 has the evidence classification for each interaction according to their “strongest” supporting experiment (Supplementary Table 1-2 and Supplementary Figure 1).

**Figure 2.**
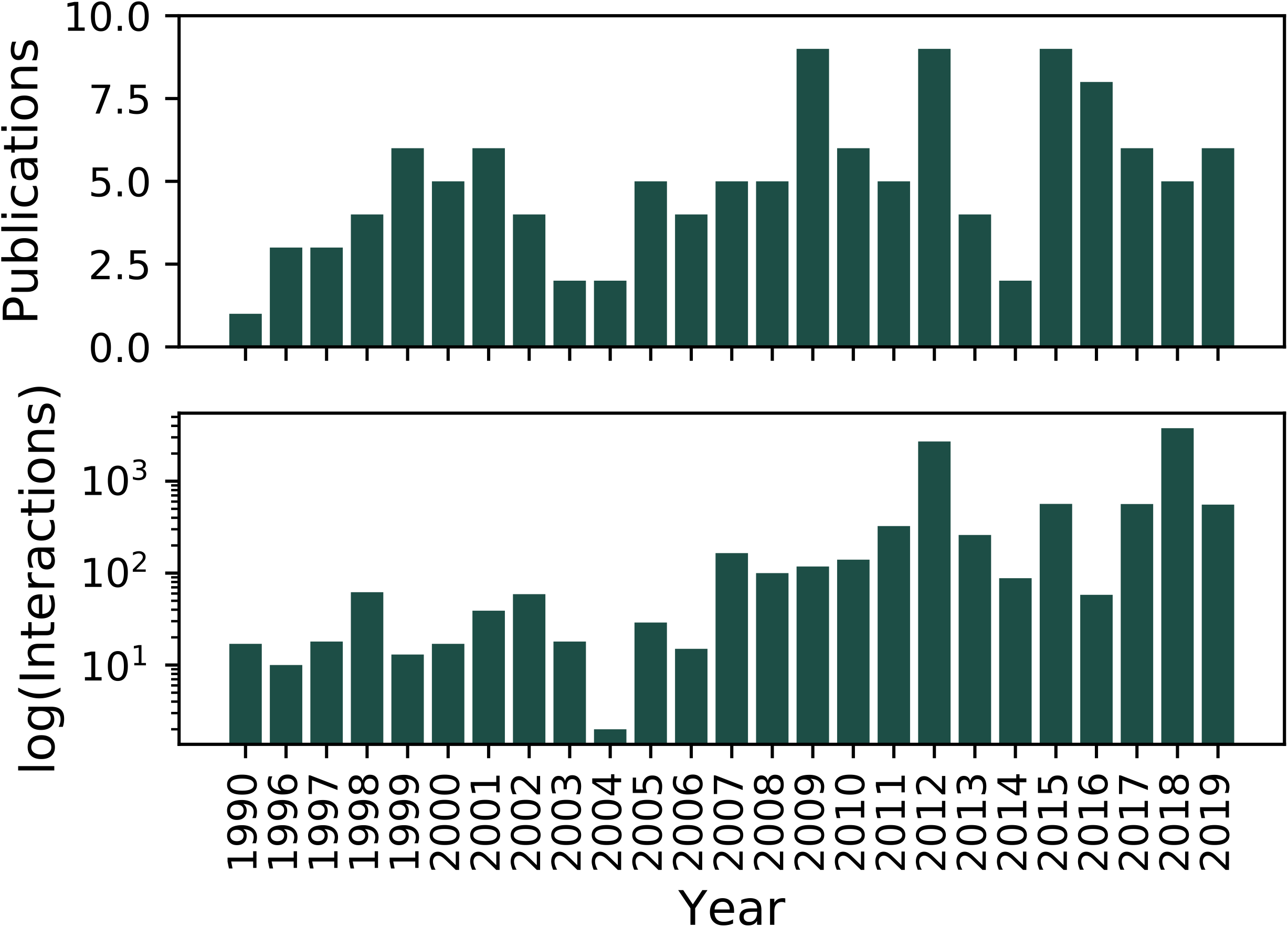
Interactions curated from literature for *Streptomyces* coelicolor A3(2). a) Number of publications per year and b) Number of interactions reported per year.

Afterward, we gathered these interactions along with others from the databases, RegTransBase^24^ available at Abasy Atlas database (https://abasy.ccg.unam.mx)^19^, and DBSCR (http://dbscr.hgc.jp/). We processed these curated interactions to construct the corresponding GRNs, removing redundancy by mapping the gene identifiers to locus tags and merging interactions while preserving the information about the effect and evidence classification of the supporting experiments, as previously reported^19^. From our curation, we reconstructed a total of seven curated networks with different evidence classification and completeness. 1) *Curated_FL* with a total of 9454 unique interactions, from which ∼5% (493/9454) are “strong”. 2) *Curated_FL*(cS) with 438 “strong” interactions from *Curated_FL*. 3) *Curated_DBSCR* with the 341 interactions from DBSCR and used the ∼34% (115/341) “strong” interaction to reconstruct 4) *Curated_DBSCR*(S). 5) *Curated_RTB* is the network from the RegTransBase database with 330 interactions, all of them labeled as “weak” since their experimental evidence was not available. Later, we merged *Curated_FL, Curated_DBSCR*, and *Curated_RTB* into 6) *Curated_FL-DBSCR-RTB* a meta-curated network comprising a total of 5386 genes and 9707 non-redundant regulatory interactions, which is the most extensive experimental GRN of *S. coelicolor* up to date. From this meta-curation, we filtered the 480 “strong” interactions to reconstruct 7) *Curated_FL(cS)-DBSCR(S)*. All curated networks are further described in Table 1.

**Table 1.**
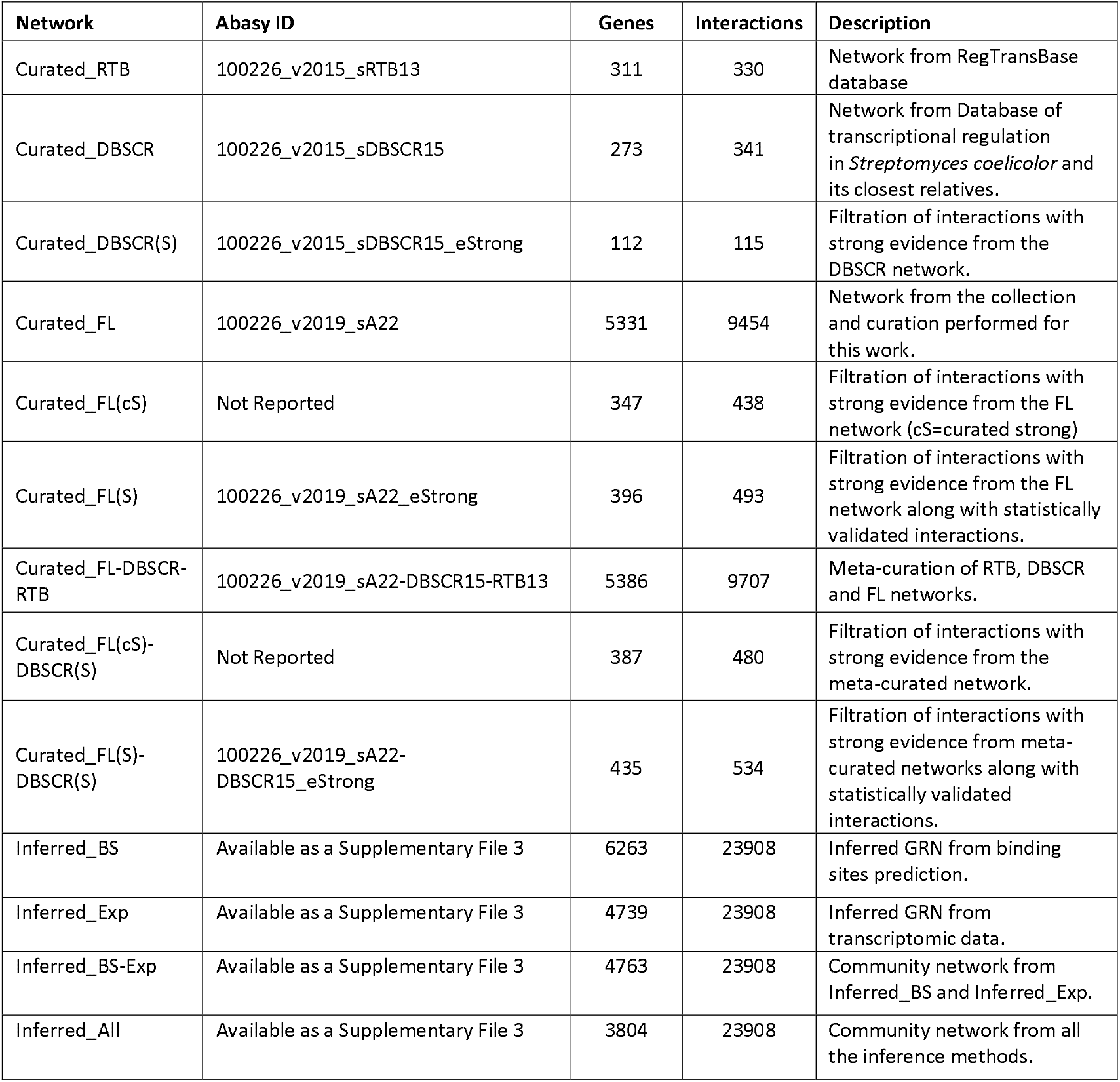
Description of networks used in this work.

### The functional architecture of the *S. coelicolor* GRN

To reveal the functional architecture and to elucidate the regulatory and biological function of some of the genes and interactions curated, we applied the Natural Decomposition Approach (NDA) on all the curated networks. The NDA is a biological-mathematical criterion to suggest a biological function of each gene based on the structure of the GRN^20^. It classifies the genes into one of the four structural classes: i) global regulators (GR), coordinating genes from different metabolic pathways^25^; ii) modular genes, group of genes working together to carry out a biological function^20,26^; iii) intermodular genes, integrating at the promoter level the response from different modules^20,27^; and iv) genes constituting the basal machinery of the cell. We decided to further study the NDA analysis of *Curated_RTB-FL-DBSCR* since it is the most complete GRN.

The NDA analysis of the meta-curated network *Curated_RTB-FL-DBSCR* revealed 20 GRs (0.37% of the 5386 network genes), 502 modular genes (9.32%), 18 intermodular genes (0.33%), and 4846 basal machinery genes (89.97%). The classification of each gene can be found at https://abasy.ccg.unam.mx/genes?regnetid=100226_v2019_sA22-DBSCR15-RTB13&class=All. Through the NDA analysis, we found 35 gene modules. From them, module number 16 is a mega-module, which is divided into 12 submodules that are connected through the intermodular genes (Supplementary Figure 2a). To analyze the GRs identified in the meta-curated network, we reviewed the literature to identify the TFs that have been previously reported as global or pleiotropic regulators in *S. coelicolor. Martín et al*.^28^ reported a detailed description of the cross-talking between the global regulators in *S. coelicolor* and other *Streptomyces*. The review provides a list of genes considered as global and wide-domain regulators, due to the hundreds of genes they regulate and the multiple effects they produce^28^. Nine out of the 20 (45%) GRs identified by the NDA were reported as such in this review. We further screened the literature to identify GRs or pleiotropic regulators reported in individual papers (Supplementary Table 4). We found 20 pleiotropic TFs or GRs reported individually, from which 13 (65%) were categorized as GRs by the NDA. See the “Global regulators” section in Supplementary File 1 for further description of the GRs identified.

This analysis also revealed 18 intermodular genes. Some of their promoters integrate the signals of different GR related to carbon, nitrogen, and phosphate metabolism. For instance, glnA (SCO2198), glnII (SCO2210), and the amtB-glnK-glnD (SCO5583-85) operon, which are known to be mediators between the nitrogen and phosphate metabolism through the binding of their GR (PhoP and GlnR) to these intermodular genes promoters ^29^. Others integrate signals from primary and secondary metabolism, or morphological differentiation and antibiotic production. A further description of these genes can be found in Supplementary File 1 in the section “Intermodular genes”. Moreover, the functional annotation of the modules identified by the NDA also provides a new functional hypothesis for genes whose function is currently unknown using a guilt-by-association strategy, as previously described^30^. From the 46 modules and submodules in the GRN, 26% (12/46) are annotated. The annotation of each module can be found at https://abasy.ccg.unam.mx/modules?regnetid=100226_v2019_sA22-DBSCR15-RTB13. Most of the annotated modules are related to cellular metabolism, organic substances metabolism, and biosynthetic processes, which are fundamental processes for every cell (Supplementary Figure 2b). We found 245 with no previous annotation in GOA^31^ assigned to the annotated modules (Supplementary Table 5).

### GRN inference based on binding sites identification performs better than that based on transcriptomics

Despite the exhaustive curation, the meta-curated network *Curated_FL-DBSCR-RTB* has regulatory information for only ∼65% (5386/7825) of the *S. coelicolor* genome (network genomic coverage) and has ∼41% (9707/23908) of its expected total interactions (network interaction coverage or completeness)^19^, considering both “strong” and “weak” interactions. We leveraged the large corpus of high-throughput data available to computationally infer missing regulatory interactions to expand our GRN reconstruction. The inference was performed from two different approaches. For the first approach, we performed a regulon reconstruction through the de novo identification of TF binding sites and linked them to downstream genes. The regulon reconstruction was based on the network *Curated_FL*(cS)-DBSCR(S) using three methods for motif discovery: MEME, Bioprospector, and MDScan (see Methods). For the second approach, we performed a GRN inference from transcriptomic data. We used seven methods for GRN inference based on the gene expression data: CLR, Friedman, GENIE3, Inferelator, MRNET, Statmodel, and TIGRESS (see Methods and Supplementary Table 6a). These methods were selected based on their performance in a previous benchmarking of GRN inference methods^18^, their availability, complete documentation, and maintenance. For the inference, we selected an Affymetrix dataset (Platform GPL9417) from NCBI GEO^33^. See the “Transcriptomic Data” section in Supplementary File 1 for further description of the data selection. Since most of the experiments in the curation were performed on the *S. coelicolor* A3(2) strain M145, which is plasmid-free, we restricted the inference to interactions among genes of the chromosome. Next, we evaluated the inferred GRNs computing the AUROC and the AUPR of the predictions (see Figure 3a and Supplementary Figure 4). From the AUPR, it is evident that in general GRN inference from binding sites performed better than the inference from expression data. For the inference from binding sites, MEME performed better than the other methods. For the inference from gene expression data, TIGRESS performed better, followed closely by GENIE3 and Inferelator. We assessed the inferred GRNs based mostly on the AUPR since it is more informative for imbalanced datasets^36^ as it is the case of GRNs inference^37^. Binding sites for each one of the interactions identified using MEME are reported in Supplementary File 3.

**Figure 3.**
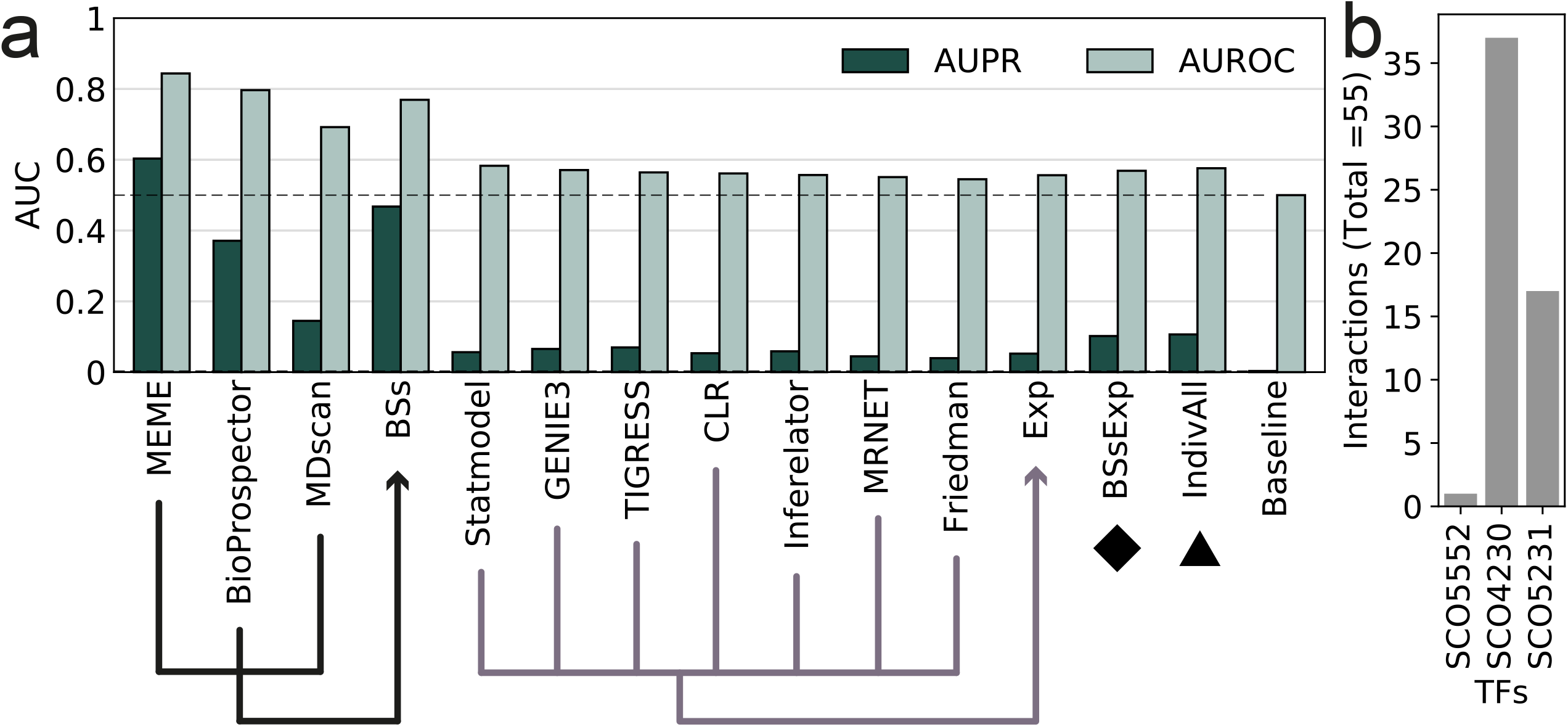
a) AUROC and AUPR for each of the methods and the community networks. b) Number of interactions statistically validated by TF

For the assessment, all the inferred networks were pruned to the 23908 best scoring interactions among genes part of the GS, since it is the number of interactions expected in the final network for *S. coelicolor*^38^. Nevertheless, the GS has only 387 interactions, which is ∼1.6% of the 23908 regulatory interactions expected in the complete regulatory network of *S. coelicolor*^19^. For this reason, the assessment only reflects the capacity of the methods to infer the interactions in the GS, while novel interactions (actual interactions not part of the incomplete GS) are labeled as false positives. Moreover, as the GS was used as prior for the regulon extension, it might provide an advantage for the network predicted by motif discovery. Because of its approach, inference by motif discovery predicts direct regulatory interactions, while inference from transcriptomic data predicts both direct and indirect ones without distinction. Thus, as the GS is only built by direct interactions, it is expected that inferred networks with the same type of interactions get a higher score. However, using the “non-strong” GRN as GS could drive to a bigger problem because indirect regulatory interactions might be spurious and are not adequate to assess causal interactions.

Given that the current GS is still quite incomplete, and we cannot do proper discrimination among the different inferred networks, instead only using the single best method, we decided to build a community network for each one of the approaches. i) *Inferred_BS* for the prediction from binding sites; ii) Inferred_Exp for the prediction from expression data; iii) *Inferred_BS-Exp*, a community network from both previous community networks; and iv) *Inferred_All*, a community network built mixing individual networks from both approaches (see Table 1). For the latter, we used the three methods for binding site inference, along with Statmodel, GENIE3, and TIGRESS from expression-based GRNs (due to their superior performance) to balance both approaches. *Inferred_BS* outperformed the rest of the community networks at both AUPR and AUROC (see Figure 3a). However, it was outperformed by MEME at both metrics. Given MEME’s outstanding performance (see Figure 3a), we used it to perform a statistical validation of “weak” interactions supported by ChIP-data, similarly as proposed in^22^ (see Methods). A total of 55 “weak” interactions were reclassified as “strong” (see Figure 3b and Supplementary Table 6b). We found one of these interactions (SCO4230-SCO4878) already reported as “strong” in the DBSCR database (*Curated_DBSCR*(S)). These statistically validated interactions were merged with the “strong” interactions from *Curated_FL* and from the meta-curated network *Curated_FL-DBSCR-RTB* into two networks: *Curated_FL(S)* and *Curated_FL(S)-DBSCR(S)*. We reassessed the network predictions with *Curated_FL(S)-DBSCR(S)* as GS and the results remained virtually the same (Supplementary Figure 5). For completeness, we also performed the assessment using *Curated_FL-DBSCR15-RTB13* as the GS (Supplementary Figure 6). However, the 72 regulators in *Curated_FL(S)-DBSCR(S)* are the only TFs that were used for the predictions based on binding sites. On the other hand, *Curated_FL-DBSCR15-RTB13* has 137 TF. This resulted in a poor recall by the predictions based on binding sites (Supplementary Figure 6), as interactions for 65 TFs are not predictable because their regulations might be carried out indirectly, with no need for a DNA binding site.

### Inferred networks have a similar structure to the largest curated networks

Even though the AUPR and AUROC metrics allow the assessment of the predictions, both metrics heavily rely on the ranking of the predicted interactions. Moreover, the GS is not complete and missing interactions would be still classified as false positives, decreasing the score more the higher their ranking is. Therefore, we assessed the inferences in terms of their structural properties and compared them against the curated networks to compensate for such drawbacks. Note that this approach has its caveats. The global structural properties of the network might be different once the GS is complete, this can be approached by comparing the predictions to all the curated networks, each of them with different completeness. Also, two networks could have the same topology with different node entities. For this reason, we use the topological assessment in complement to the AUPR and AUROC metrics to identify the best prediction.

One of the main structural properties of biological networks is that they are scale-free and hierarchically modular. Same properties that our curated networks have been proved to possess (Supplementary File 1). Therefore, as an initial approach, we asked whether the inferred networks are scale-free too. The degree (nodes’ connectivity) distribution *P*(*k*) of scale-free networks follows a power law, *P*(*k*) ∼ *k*^-*α*^, with 2 < *α* < 3 ^20,38,39^. If *α* = 2, there is a unique global regulator (hub-and-spoke network) and if *α* > 3, scale-free networks lose most of their characteristic properties^39^. First, to compute this α for the inferred networks, we performed a robust linear regression over a log-log plot of the complementary cumulative degree distribution and corrected the exponent accordingly (see Table 2 and Supplementary Figures 7-11). All inferred networks’ degree distribution seems to follow a power law according to the adjusted coefficient of determination. Nevertheless, the data points in *Inferred_All* appear to be divided into three regions with different tendencies, instead of the two that are present in the other networks (Supplementary Figures 7-11). Usually, this type of network is divided into two regions, the region of the nodes (genes) with a low degree, and the one with nodes with a high degree^40^. The appearance of a third region might be a consequence of merging networks from methods with different approaches. This could affect the structural properties of the merged networks, while communities from the same approach appear to have more similar structural properties. In the case of *Inferred_BS-Exp*, the construction of communities ahead by each approach create more compatible networks in terms of structure that can be conveniently mixed.

**Table 2.** Network properties for inferred networks

| Property | Inferred_BS | Inferred_Exp | Inferred_BS-Exp | Inferred_All | Curated_FL- |
| --- | --- | --- | --- | --- | --- |

|  |  |  |  |  | **DBSCR-RTB** |
| --- | --- | --- | --- | --- | --- |
| Number of nodes (N) | 6263 | 4739 | 4763 | 3804 | 5386 |
| $\ln(\ln(N))$ | 2.17 | 2.14 | 2.14 | 2.11 | 2.15 |
| Average shortest path length | 2.86 | 3.38 | 3.38 | 3.11 | 2.84 |
| Average clustering coefficient | 0.213 | 0.385 | 0.385 | 0.470 | 0.182 |
| $\alpha (P(k))$ | 1.861 | 1.952 | 1.955 | 1.968 | 1.742 |
| $R^2_{adj}$ | 0.87 | 0.92 | 0.92 | 0.91 | 0.84 |
| $\alpha (C(k))$ | 0.924 | 0.767 | 0.742 | 0.729 | 1.142 |
| $R^2_{adj}$ | 0.79 | 0.58 | 0.54 | 0.68 | 0.89 |

To confirm that their degree distributions follow a power law, we contrasted each distribution of the inferred networks against alternative fat-tailed probability distributions (Power-law, exponential, stretched exponential, lognormal, and truncated power-law) using Kolmogorov– Smirnov tests^38,41^(Supplementary Table 6a). We found that the degree distribution of the inferred networks adjusted better to a power-law than to an alternative distribution. Then, we computed a maximum likelihood estimation for the exponent (α) of the power laws and found that most of them are between two and three, except for *Inferred_All*. This shows it as an anomalous scale-free network (Supplementary Table 7). Perhaps caused by the mixing of networks with diverse structural properties. Nevertheless, we could consider all inferred networks to be scale-free.

Furthermore, we checked other properties of scale-free networks (see Table 2)^39,42^. The four community networks have small average shortest path lengths and a high clustering coefficient. *Inferred_BS* has the smallest average short path length, while *Inferred_All* has the highest average clustering coefficient (see Table 2). Scale-free networks also present an ultra-small world effect, which implies that the average path length is proportional to ln(ln(*N*)) (N is the number of nodes in the network). This is the case for all the inferred networks. Another characteristic of GRNs is their hierarchical modularity, which implies a diamond-shaped hierarchical organization as has been revealed by the NDA^20^. In a scale-free network, this implies that the clustering coefficient depending on the degree (*C*) follows a power law as *C*(*k*)∼*k*^-139^. *Inferred_BS* has the exponent closest to -1 (−0.92), with the best *R*^2^. Even though *Inferred_BS* seems to be the network that behaves closest to a GRN, all networks have similar values, which makes it difficult to discern the most reliable inferred network by this approach.

We included several other structural properties of the networks to perform a more thorough comparison (Supplementary Table 8). We clustered the vectors of structural properties for the curated and community-inferred networks (see Figure 4a). The clustering partitions the networks into two major groups. The first one contains the curated networks and *Inferred_BS*, while the second group contains the other inferred networks. The first group is in turn also divided into two groups: one with the two largest curated networks and *Inferred_BS*, and the other one with the remaining curated networks. The reason for this may be due to the size of the networks (see Table 1). When standardizing the property values by max-min feature scaling, the two largest curated networks were clustered with the predictions (Supplementary Figure 12), which could be also due to the size of the network. To reduce the network size influence we used the network dissimilarity measure proposed by Schieber *et al*.^43^. We considered the third term which makes the distance measure robust to graph size in terms of the number of nodes (genes)^43^ (see Figure 4b). Even with this metric, the two largest curated networks were clustered with the inferred networks. This might be a consequence of the high value of maximum out-connectivity and structural genes in the largest curated networks, like those found in the inferred networks. This shows that the inferred networks are similar that the most complete curated networks in terms of structure, suggesting their reliability.

**Figure 4.**
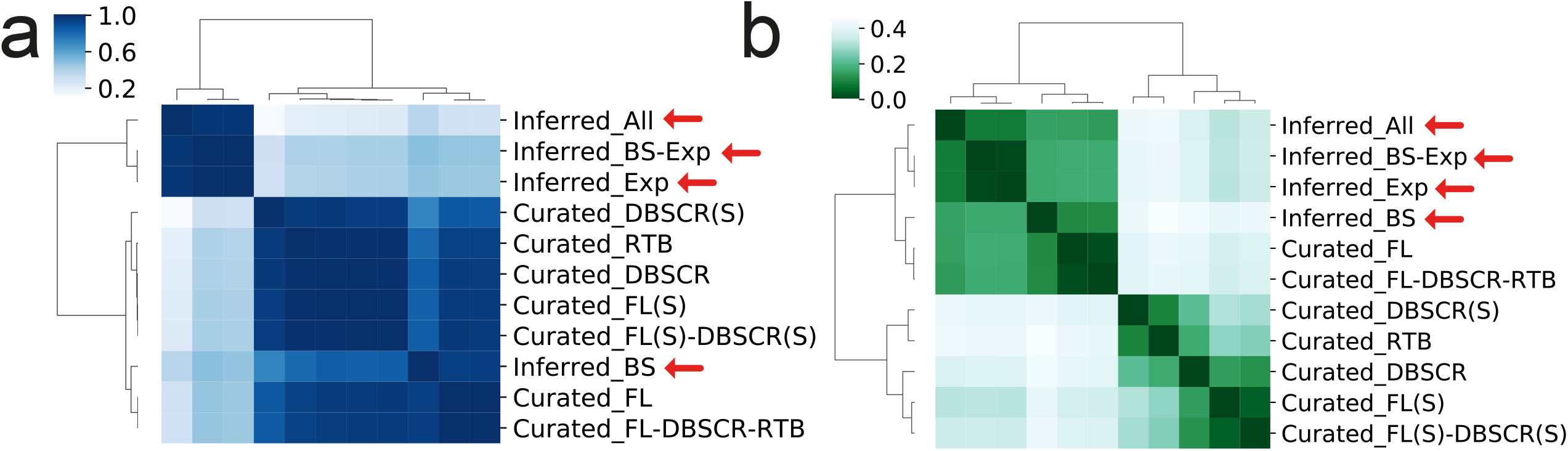
Network comparative by structural properties. a) Pearson correlation of the profile of structural properties listed in Supplementary Figure 12. b) D-value from Schieber *et al*.^43^ to measure network similarity.

### The Natural Decomposition Approach identifies regulatory networks similarity despite their different completeness level

We compared all the curated and community inferred networks based on the Simpson similarity index of the four components proposed by the NDA: global regulators, modular genes, intermodular genes, and the basal machinery (see Figure 5). The Simpson similarity index measures the size of the core of two sets with reference to the smallest one^44^. Note that two identical sets, being one a subset of the other will have a score of 1. On the other hand, two completely different sets will have a score of 0.

**Figure 5.**
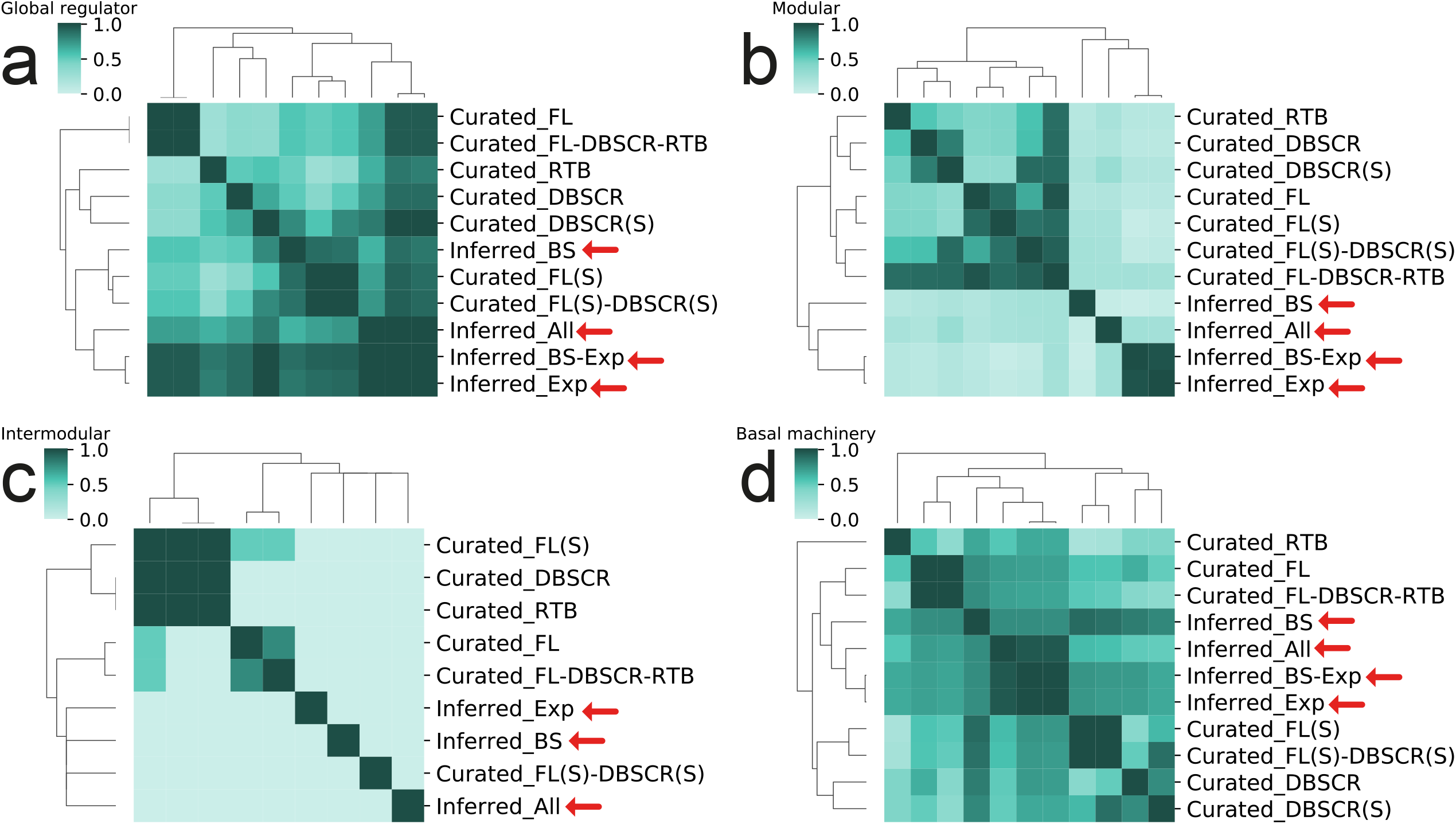
Simpson similarity index for NDA analysis for all curated and inferred networks. a) Global regulators. b) Modular genes. c) Intermodular genes. d) Basal machinery genes.

When comparing the global regulators (see Figure 5a), there is not a distinct division among the networks. *Infered_BS-Exp and Infered_Exp* have a similar correlation with all the curated networks, slightly higher with *Curated_DBSCR(S), Curated_FL*, and *Curated_FL-DBSCR-RTB*. These two inferred networks have the highest amount of GR, 116, and 114 respectively; thus, the other GRs predicted could be easily a subset of them. For the case of *Infered_BS*, it has the highest correlation with the “strong” networks. This is expected since these networks have only direct regulatory interactions, as the interactions predicted in *Inferred_BS*, while there is no evidence of direct regulation for transcriptomic-based inferred interactions. This can result in an underestimation of the effect of GRs over the rest of the genes since GRs which regulate several targets from different processes might not be predicted as such in the “strong” networks. However, when their indirect influence is represented in the network, their ranking as GRs is noticeable.

When analyzing the modular genes, there are two major groups (see Figure 5b): the major group with the curated networks is divided into two subgroups, one contains *Curated_RTB, Curated_DBSCR*, and its “strong” version *Curated_DBSCR(S)*, and the second subgroup contains the integrations and curations proposed in this work. Interestingly, the meta-curated network *Curated_FL-DBSCR-RTB* correlates very well with all the smaller networks it contains, from which we could deduce that modular genes are conserved despite the addition of new regulatory interactions. In the second group, composed of the inferred networks, we can see there is not a high correlation among them. *Inferred_BS* is the closest to the curated networks, while Inferred_Exp and *Inferred_BS-Exp* have a high correlation. This tells us that the interactions in Inferred_Exp have a larger influence on the module configuration of *Inferred_BS-Exp* than *Inferred_BS*. The difference between the curated and inferred networks might come from the fact that inferred networks have a greater number of GRs and a much lower number of modular genes when compared with the curated networks (Supplementary Table 9).

Intermodular genes are the less conserved NDA class (see Figure 5c). There is overlap only among the smallest curated networks, all share the intermodular gene SCO5877, which appears as a TF in the other curated networks. Moreover, there is an overlap between *Curated_FL* and *Curated_FL-DBSCR-RTB*, which share most of the interactions. Thus, is expected that they also share most of the intermodular genes. Note that the networks *Curated_DBSCR(S)* and *Inferred_BS-Exp* are not included in the clustering since they did not present any intermodular genes. Finally, when analyzing the basal machinery (see Figure 5d), the larger curated networks are grouped on one side, next to the inferred networks, and finally the smallest curated networks with *Curated_RTB* as an outgroup. Even though *Inferred_BS* is grouped with the other inferred networks, it has a higher correlation with *Curated_FL(S)* and *Curated_FL(S)-DBSCR(S)*, which again evidence the similarity among these three networks.

### Assessment of the global regulators’ inference

In the NDA, the identification of global regulators is a key step in the classification of every node in the GRN. Previously, it has been reported a high overlap between the predictions of global regulators by the NDA and those reported in the literature for *E. coli*^20^, *B. subtilis*^45^, and *C. glutamicum*^27^. We used the set of GRs reported by *Martín et. Al*.^28^, besides those reported in independent articles, and the union of both sets (Supplementary Table 4). Then we assessed the predictions of the GR using the Matthews correlation coefficient (MCC) (see Figure 6) We used the MCC score as it is more informative and reliable than F1 for binary classification assessment^46^, however when applying the F1 score we obtained consistent results (Supplementary Figure 13).

**Figure 6.**
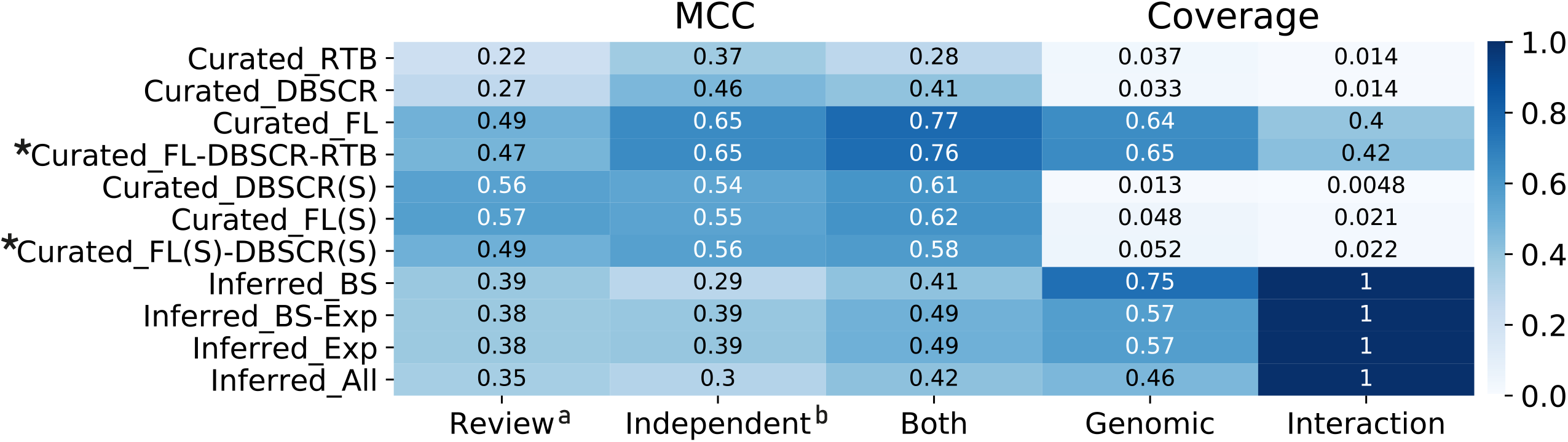
MCC for global regulators predicted by NDA for each of the curated and inferred networks. Scores ≥ 0.5 are represented in white numbers and meta-curations are marked with an asterisk. ^a^Gold standard curated from reference ^28^. ^b^Gold standard curated from independent publications (Supplementary Table 4).

*Curated_FL* and *Curated_RTB-FL-DBSCR* have the best performance in GR prediction. However, the “strong” networks have a slightly smaller score even having much less genomic coverage. This shows that the GRs are very robust to perturbations in the network as previously shown^20^. On the other hand, despite the high coverage of the inferred networks, the performance of the predictions with such networks was poor. This could be, as it was mentioned before, due to the great amount of GR predicted by these networks, which would cause a high proportion of false positives affecting the score. *Inferred_BS* produced the most conservative prediction (lowest false-positives rate) among the inferred network (Supplementary Figure 14 and Supplementary Table 9).

### GRN inference from transcription factor binding sites proves to be the most reliable approach and allows the prediction of new TFs for the most studied SARPs

*Inferred_BS* performed the best on AUPR and AUROC among the community inferences and is the most similar to the curated networks according to its structural properties and system-level components. Moreover, it has the largest genomic coverage among all the networks, which would be advantageous for a deeper study of transcriptional regulation in *S. coelicolor*. Therefore, we considered *Inferred_BS* as the most reliable inferred network, despite that similar studies suggest the integration of inference approaches as the most suitable methodology for GRN reconstruction^18,47,48^.

Thus, we decided to use *Inferred_BS* to further study the regulation of the SARPs of the most studied antibiotics in *S. coelicolor*: ActII–orf4, RedD/RedZ, CpkO (also known as KasO), and CdaR, which regulate the production of ACT, RED, yCPK, and CAD, respectively^3^ (Supplementary Table 10). A total of 13 new putative regulators for the SARPs were predicted, providing a great opportunity to find new targets to manipulate the *S. coelicolor* antibiotic production. As it is not possible to computationally confirm a direct binding of this regulator, it is necessary to corroborate them with wet-lab experiments.

Following, we describe some of the TFs predicted for the SARPs using the workflow suggested in Figure 1b: For *actII-orf4* (SCO5085) only one novel regulator was inferred, MacR (SCO2120) which is the response regulator of the TCS MacRS, while the rest was already part of Curated_FS(S)-DBSCR(S) (Supplementary Figure 15). This TCS has been proved to activate ACT production. Nevertheless, a ChIP-qPCR analysis was not able to prove an *in vivo* interaction between MacR and *actII-orf4*, although a direct interaction was not tested^49^. For *redD* (SCO5877) two new regulators were predicted, LipR (SCO0712) and ActII-orf4 (SCO5085). LipR is related to AfsR (SCO4426)^50^, homolog to the SARPs, and activator of the ACT and RED production^51^. Moreover, its mutant affects ACT production^50^, which makes it plausible to affect RED production as well. It has been suggested that ActII-orf4 might regulate the production of other antibiotics^4^, which could be by binding directly to their CSR. For *redZ* (SCO5881) five new regulators were predicted, among them is GluR (SCO5778) which has been shown to affect RED production. Nevertheless, it has been shown that GluR does not bind directly to *redZ*, thus it could an indirect regulation^52^. Another one, StgR (SCO2964) has been shown by an RT-qPCR experiment to be a repressor of *redD*^53^. This repression could be through the direct binding to *redZ*. HpdA (SCO2928) and HpdR (SCO2935) are related to tyrosine catabolism, which produces important precursors for antibiotic biosynthesis^54^. Moreover, HpdA has been shown to activate actII-ORF4, therefore might have a more direct role in RED production. In the case of cdaR (SCO3217), we have four predicted regulators, among them, *OsdR* (SCO0204) and RamR (SCO6685). Both are related to the response to stress and the development of *S. coelicolor*^55,56^. SsgR (SCO3925) regulates sporulation and morphological differentiation^57^. These all processes are highly related to antibiotic production. Finally, for *cpkO/kasO* (SCO6280) six new regulators were inferred, among them *OsdR* (SCO0204), LipR (SCO0712), and StgR (SCO2964) were described before. Another one is NnaR (SCO2958), which regulates spore formation and antibiotic production^58^. We refer the reader to the Supplementary material for the complete list of the predicted regulators in every predicted network (Supplementary Table 10).

We further compared the six regulators of *actII-orf4* identified by *Inferred_BS* with the 11 TFs found in *Inferred_All*. From the latter, only two are part of *Curated_FS(S)-DBSCR(S)*, and three more are included in *Curated_FS-DBSCR-RTB* (Supplementary Figure 15,16). We found that the interactions included in *Inferred_All* tend to be carried out indirectly (i.e., not through a TF-DNA interaction), suggested by a poor overlap between *Inferred_BS* and *Inferred_All*, and biological insights such as the putative regulation of *actII-orf4* by *OsdR*.

### Comparative analysis with *Corynebacterium glutamicum* shows coherent system-level components conservation

The diamond-shaped structure identified by the NDA is conserved between E. coli and B. subtilis^45^. As an application of the meta-curated network, we studied the conservation of its system-level components, comparing it against the *C. glutamicum network. C. glutamicum* is phylogenetically related to *S. coelicolor*, and a model organism for the study of GRNs^27^. We applied the regulogs analysis^59^ with one-to-one orthology relationships to alleviate network incompleteness and make them comparable. As prior networks, we used *Curated_FL(S)-DBSCR(S)* (534 interactions) for *S. coelicolor* and 196627_v2020_s21_eStrong from Abasy Atlas^19^ (2941 interactions) for *C. glutamicum*^16^. After the regulogs analysis, we ended up with 2966 interactions in *C. glutamicum* and 692 interactions in *S. coelicolor*.

We used the complemented networks to identify GRN-wide orthologous relationships defined as the orthologous present in the GRN of the respective organism. We obtained a total of 188 GRN-wide orthologous relationships from a total of 995 1:1 orthologs identified by OrthoFinder^60^. We applied the NDA analysis to both GRNs to identify the system-level components and computed the fraction of the GRN-wide orthologous in each combinatory relationship between the NDA classes (see Figure 7a). We found that most of the GRN-wide orthologous (54%) are classified as basal machinery in both organisms. This is expected since 73% and 74% of the genes correspond to the basal machinery in the complemented networks of *C. glutamicum* and *S. coelicolor*, respectively. Besides, the distribution of the genes in the chromosome of *S. coelicolor* shows a central core, where are genes likely related to primary functions such as DNA replication, transcription, translation, and amino-acid biosynthesis; and likely non-essential genes such as secondary metabolism are in the chromosome arms^5^. More than 59% (111/188) of the GRN-wide orthologs conserved the same class in both organisms (Figure 7a) showing high conservation of the NDA classification.

**Figure 7.**
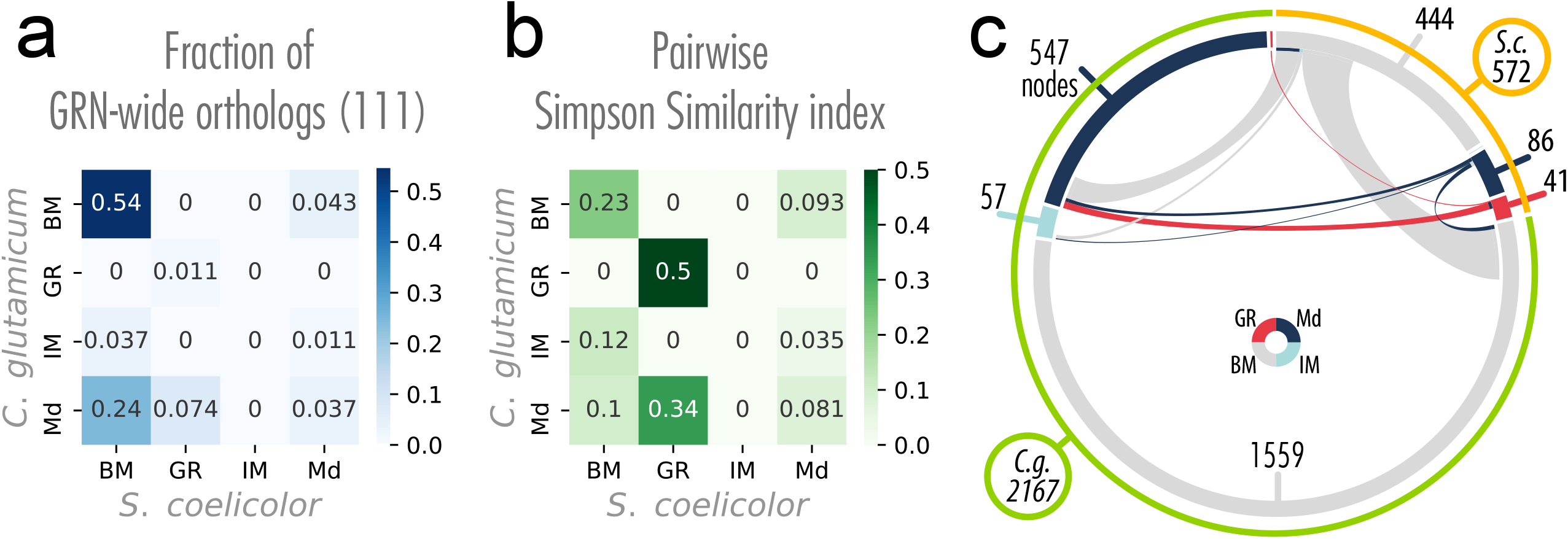
Conservation of the systems-level components between *S. coelicolor* and *C. glutamicum*. a) NDA classification for the GRN-wide orthologs in their corresponding organism. Total matrix sum to 1. Most of the GRN-wide orthologs are classified as basal machinery in both organisms b) Overlapping of the NDA classes between the two networks with reference to the smallest set. Each cell can range between 0 and 1, where 1 means one class is a subset of another, and 0 means there is no overlap at all. c) Similarities between the two organisms highlight the size difference between the datasets. The color of the inner circle sections represents the NDA classes and ribbons colors represent the *S. coelicolor* NDA classes. Numbers represent the genes for each class and organism. Gray ribbons are the widest ones, representing that all the basal machinery of *S. coelicolor* with 1:1 orthology relationship with *C. glutamicum* is classified either as basal machinery or modular genes in *C. glutamicum*. This suggests that multiple basal machinery genes could be reclassified as modular components in more complete reconstructions of the *S. coelicolor* network.

We studied the pairwise Simpson similarity index between the four classes between the two organisms to remove the problem of the imbalanced classes (see Figure 7b). GR is the class with the highest conservation rate, the orthologs of seven of the eight GRs in *C. glutamicum* are also GRs in *S. coelicolor* (see Figure 7b and Figure 7c). The conservation between the same class in the two organisms is also high for the basal machinery, while poor for the modular genes. For the case of intermodular genes, even though the networks were complemented with information from the other network, they are not conserved at all (see Figure 7b). Previous work reported intermodular genes as the least conserved of the system-level components^27^. Intermodular genes are the most likely responsible for giving the GRN flexibility and increasing evolvability by scouting different combinations of regulatory interactions between physiological functions so the organism could adapt better to environmental changes^45^. These results agree with a previous analysis of the robustness of the NDA to a random node and edge remotion showing GR and intermodular genes as the most and least conserved classes, respectively^27^.

On the other hand, 24% of the GRN-wide orthologs that are modular genes in *C. glutamicum* were classified as basal machinery in *S. coelicolor*. This could be due to three possible reasons^45^: i) the basal machinery genes in *S. coelicolor* are misclassified and further research is needed to find the missing regulatory interactions (see Figure 7c) that will integrate some of these genes into a module. ii) The GRs controlling *C. glutamicum* genes are not yet identified as GRs. iii) Genes in *S. coelicolor* need a more direct regulation because of their physiological function (high plasticity of transcriptional regulation). A previous genomic comparison between *S. coelicolor, Mycobacterium tuberculosis*, and *Corynebacterium diphtheriae* showed a synteny among the whole chromosome of these last two microorganisms and the core of the one of *S. coelicolor*^5^. *C. glutamicum* is phylogenetically closely related to M. tuberculosis and *C. diphtheriae*, with roughly similar genome size. Therefore, a similar result would be expected. Furthermore, as more classical experiment data become available, new regulations for the currently basal machinery would turn those genes into the modular class. However, a deeper analysis of diverse factors such as genome size, the niche of the organisms, and a wider range of organisms are required to further study the robustness of the NDA analysis.

## Conclusions

A meta-curated regulatory network for *S. coelicolor* (*Curated_RTB-FL-DBSCR*) was reconstructed from a collection and curation of regulatory interactions experiments in literature and databases. From the NDA analysis of the meta-curated network, we could identify 20 global regulators, of which 95% (19/20) have already been reported as global or pleiotropic regulators. 46 functional modules were identified along with 18 intermodular genes, some of them found to be involved in more than one biological process. Functional modules annotated by GO enrichment allowed via a ‘guilt by association’ strategy to propose a function for 245 genes without any previous functional annotation or annotated as ‘hypothetical protein’. This network is, however ∼42% of the estimated complete regulatory network, which evidences a lack of information related to *S. coelicolor* transcriptional regulation. Especially for interactions experimentally supported by “strong” evidence, which accounts for only ∼2% of the estimated complete network. We indeed found a low level of direct experimental validation for the regulatory interactions reported in the literature and curated in this work as only ∼6% (533/9687) are supported by experiments confirming the binding of the TF to the upstream region of the target gene, the so-called “strong” evidence. The low level of “strong” evidence is due to the high fraction of high-throughput experiments aimed to unveil the regulatory network. This highlights the importance of carrying on classical experiments aimed to confirm the weakly supported interactions (e.g., EMSA, *in vitro* transcription assay, and DNA footprinting) to increase our knowledge of the transcriptional regulation in *S. coelicolor*. Notwithstanding, the meta-curated network *Curated_RTB-FL-DBSCR* provides the most extensive and up-to-date reconstruction available for the regulatory circuitry in this organism and already portrays accurately the functional organization of *S. coelicolor* regulation. GRN inference from transcriptomic and DNA sequence data was performed and the inference from TF binding sites identification showed to be the best approach according to interactions inference assessment, topological assessment, and systems-level comparison. This final inferred network is a valuable guide for wet-lab experiments, since narrows down the search space of the possible TF for each gene. Besides, it can be used in computational models of *S. coelicolor*. From this network 13 new TFs were predicted to bind in the upstream region of five of the principal SARPs, most of which previously proved to affect indirectly antibiotic production or to be related to stress response or morphological differentiation. Finally, we compared *S. coelicolor* network to *C. glutamicum* GRN, showing one of the many potential applications of the curated network. There we found high conservation only for the basal machinery, which might be a result of the high plasticity of the transcriptional regulation. To visually explore the interactions validated by experiments and identify the role of the genes in the global regulatory network (e.g., global/local regulators) and functional annotation, we strongly suggest using Abasy Atlas.

## Material and methods

### Transcriptional regulatory interactions curation

We performed a comprehensive review of the literature to identify experimentally-supported transcriptional regulatory interactions in *Streptomyces* coelicolor A3(2). We searched peer-reviewed articles in Google Scholar and PubMed using the keywords “*Streptomyces coelicolor*” AND “transcriptional” and its variations AND “regulation” and its variations. In the case where reviews were found, their references were followed to the original research papers. Then, we performed the curation and organized the interactions (Supplementary File 1). Experiments were classified according to their methodology and their names were standardized for the sake of clarity and easier evidence classification. We merged these interactions with two previously curated networks, one was reconstructed from an XML provided by the DBSCR team and the other one from RegTransBase^24^ available at the Abasy Atlas website. These datasets are available from http://dbscr.hgc.jp/ and https://abasy.ccg.unam.mx. Abasy Atlas is a database of meta-curated bacterial GRNs for nine species including *S. coelicolor*^19^. It also provides historical snapshots for other model organisms such as Escherichia coli, Bacillus subtilis, *Corynebacterium glutamicum*, and *Mycobacterium tuberculosis*^19^.

### GRN inference from transcription factor binding sites

To extend the regulons for the TFs identified in the literature, we used the set of “strong” interactions as prior (*Curated_FL*(cS)-DBSCR(S)). We reconstructed a position weight matrix (PWM) for every TF in the “strong” network using the non-overlapping up to -300 to +50 bp (with reference to the translation start codon) upstream regions of their TGs as input for three motif discovery algorithms. Namely, i) MEME, an extension of the expectation-maximization algorithm for fitting finite mixture models^61^; ii) BioProspector, based on multiple Gibbs sampling^62^; and iii) MDscan, that employs a heuristic word-enumeration approach combined with statistical modeling^63^. Then, we used FIMO^64^ (p-value threshold = 1×10^−4^) to identify TF-TG interactions. As most of the interactions curated are from *S. coelicolor* A3(2) strain M145 (plasmid-free), we excluded genes that are not part of the chromosome.

### GRN inference from transcriptomic data

We downloaded first the transcriptomic dataset for *S. coelicolor* available at the COLOMBOS database ^32^. Then, we also download data from the NCBI Gene Expression Omnibus (GEO)^33^. From there we download an Affymetrix dataset (Platform GPL9417) and an RNA-Seq dataset (GPL26763. Afterward, we normalize the Affymetrix data using Robust Multi-chip Averaging (RMA) with the affy package^65^ and used the gPCA package^66^ to identify a batch effect in the data, which was corrected with Combat from the sva package^67^, all of them are packages for R. The data counts on 137 transcriptomes for 7738 genes. As in the case of GRN inference from transcription factor binding sites, we only considered genes from the chromosome. We selected the best inference methods according to their outstanding performance in the DREAM challenge^18^. Moreover, we selected methods that have an implementation in R or Matlab and were well documented. The inference methods selected were: i) CLR^68^, a method that applies mutual information; ii) GENIE3^69^, which applies tree-based regression and feature selection; iii) Inferelator^70^, which applies regression and variable selection; iv) MRNET^71^, which applies the maximum relevance/minimum redundancy algorithm; and v) TIGRESS^72^ which applies LARS combined with stability selection. Along with these methods, we used two modifications we propose in this work, Friedman, and Statmodel (Supplementary File 1). We provided all methods with a list of 137 TFs from the meta-curated network *Curated_FL-DBSCR-RTB* to infer causality^18,72^.

### Integration of individual inferences into a community GRN

To increase the precision of the predictions we used a community approach^18^ integrating individual predictions from different algorithms. First, the individual predictions are sorted by their confidence score, keeping the most reliable ones at the beginning of the prediction list. Then, the average of the rank positions in the predictions is given as the community score for each interaction. For missing interactions in a prediction list, the position is equal to the size of the prediction list + 1. All community networks were pruned to the 23908 first interactions with the highest score, which is the predicted size of the complete GRN of *S. coelicolor* reported by Abasy Atlas v2.4^19^ according to the model developed in^38^. The model is constantly being updated on the Abasy Atlas website by the addition of new networks and interactions, which will cause a slight variation in the number of interactions^19^.

### Assessment of the inferred GRN

We computed the area under the precision-recall curve (AUPR) and the area under the receiver operating characteristics curve (AUROC) to assess our predictions using in-house scripts. The AUPR depicts the precision (1) as a function of the recall (2) obtained by the predictor. The AUROC depicts the relation between the recall, also called the true positive rate (TPR) (2), and the false positive rate (FPR) (3). Note that unknown actual interactions between genes in the GS will still be considered as FP^37^. For this reason, interactions involving genes that are not part of the GS were not considered.

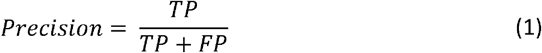

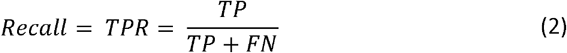

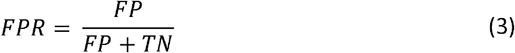

### Statistical validation for interactions supported by ChIP experiments

We performed a statistical validation approach for ChIP-data^22^ for those interactions supported by ChIP experiments and no “strong” experiment. Because of the lack of data in some articles, instead of performing the statistical validation finding the motifs from the ChIP data directly as in^22^, we used the “strong” interactions as seed to construct the matrix models for the TFs. We used MEME^61^ to build a position weight matrix (PWM) for each TF with at least 3 “strong” interactions. Then, we used FIMO with the matrix for each TF to scan the non-overlapping up to -300 to +50 bp upstream regions with reference to the translation start sites of *S. coelicolor*. We kept those TF binding sites with p-value < 1×10^−4^ and used these interactions to compare them with those supported by ChIP technologies, keeping the intersection of both sets (interactions supported by both ChIP and motif finding approaches).

### Network similarity

We computed the characteristic structural properties for GRNs reported as global properties on the Abasy Atlas database^19^. Namely, regulators (*k*_out_ > 0) (%), direct regulatory interactions, self-regulation (%), maximum out-connectivity (%), network density, weakly connected components, genes in the giant component (%), feedforward circuits, complex feedforward circuits, 3-Feedback loops, average shortest path length, network diameter, average clustering coefficient, adjusted coefficient of determination 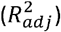 of *P*(*k*), and 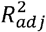of *C*(*k*). Then we used pairwise Pearson correlation among the profiles of the structural properties of the networks and cluster them according to the Euclidean distance among the correlations using Ward’s method. To compute the pairwise network dissimilarity we used a Python implementation adapted from that proposed by *Schieber et al*.^43^. We use it with the parameters proposed by the authors (0.45, 0.45, 0.1), being the last one required to discriminate network size in terms of nodes (genes)^43^. Then, we clustered the networks by using Euclidean distances among its dissimilarity values using Ward’s variance minimization algorithm as the linkage method.

### System-level components

We applied the Natural Decomposition Approach (NDA), a biological-mathematical criterion to identify the components of the diamond-shaped structure of the GRNs^20^, on every curated and inferred network. See the Supplementary methods (Supplementary File 1) for a brief description of the NDA, and^20^ for further details. The assessment of the prediction of GRs was performed using in-house scripts for MCC, precision, and F1-score. Scores were computed for each GRs prediction obtained by the NDA with the different networks analyzed in this work. As GS we used the GRs previously reported in the literature from a review^28^, from individual publications, and both (Supplementary Table 4).

### Comparative analysis against *C. glutamicum*

For the regulogs analysis, we used as prior the strong regulatory networks *Curated_FL(S)-DBSCR(S)* for *S. coelicolor*, and 196627_v2020_s21_eStrong from the Abasy Atlas database for *C. glutamicum*^19^, considering only the interactions between two genes both mapping to a locus tag. We used MEME to construct a PWM for every TF with at least three TGs using their upstream sequences. These sequences were defined as the non-overlapping regions of up to -300 to +50 bp with reference to the translation start codon and were obtained with retrieve-seq from RSAT^73^. Then, we used FIMO with the PWM of the TFs from *S. coelicolor* to find individual occurrences with a p-value < 1×10^−4^ in the upstream sequences of *C. glutamicum*. The same was done in the opposite direction. With this, we seek to alleviate network incompleteness by extrapolating known interactions from an organism to the other^59^. Predicted interactions were sorted by p-value and only the best scoring result was conserved for redundant interactions. Afterward, we used Orthofinder to find one-to-one ortholog relationships between both organisms. We used it due to its high accuracy^60^. The orthologs were used to further filter FIMO predictions to conserve interactions in which both TF and TG have a one-to-one orthologous relationship in the other organism. We considered the original “strong” network interactions at the beginning of the interactions list. The NDA was applied to both expanded GRNs to identify ortholog systems and only the genes with one-to-one orthologs in the other organism’s network (GRN-wide orthologs) were considered in the analysis.

## Availability of supporting data

The data set(s) supporting the results of this article are included within the article, its additional Files, or in Abasy Atlas at https://abasy.ccg.unam.mx/.

## Acknowledgments

The authors thank Yuko Makita and Prof. Kenta Nakai from the RIKEN Center for Sustainable Resource Science for kindly providing data from DBSCR. This work was supported by the Programa de Apoyo a Proyectos de Investigación e Innovación Tecnológica (PAPIIT-UNAM) [IN205918 and IN202421 to J.A.F.-G.]. A.Z.-A. received a doctoral scholarship [call 647 (2014)] from the Colombian Administrative Department of Science, Technology, and Innovation (COLCIENCIAS). J.M.E.-R. is a doctoral student from Programa de Doctorado en Ciencias Biomédicas, Universidad Nacional Autónoma de México (UNAM); he received fellowship 959406 from CONACYT. We thank one reviewer for helpful and encouraging suggestions.

## Authors’ contributions

Conceptualization, J.A.F.G.; Methodology, A.Z.A., J.M.E.R., J.K.G.K., and J.A.F.G.; Software, A.Z.A., J.M.E.R., J.K.G.K. and J.A.F.G; Validation, A.Z.A., J.M.E.R., and J.A.F.G; Formal Analysis, A.Z.A., J.M.E.R., J.K.G.K., and J.A.F.G; Investigation, A.Z.A., J.M.E.R., J.K.G.K., and J.A.F.G; Resources, J.A.F.G.; Data Curation, A.Z.A.; Writing – Original Draft Preparation, A.Z.A., J.M.E.R, and J.K.G.K.; Writing – Review & Editing, A.Z.A., J.M.E.R, and J.A.F.G; Visualization, J.M.E.R.; Supervision, J.A.F.G.; Project Administration, J.A.F.G.; Funding Acquisition, J.A.F.G.

## Additional Information Competing interests

The authors declare that they have no competing interests.

## Supplementary Information

- Supplementary File 1.docx: Word file with the Supplementary figures 1 – 14. NDA and structural analysis of the *Curated_FL-DBSCR-RTB* network. Supplementary methods and references.
- Supplementary File 2.xlsx: Excel file with Supplementary tables 1 – 10.
- Supplementary File 3.xlsx: Compressed folder with all the flat Files of the inferred networks.

