## Supplementary file 1 for "Curation, inference, and assessment of a globally reconstructed gene regulatory network for *Streptomyces coelicolor*"

^1^Regulatory Systems Biology Research Group, Laboratory of Systems and Synthetic Biology, and ^2^Undergraduate Program in Genomic Sciences, Center for Genomics Sciences, Universidad Nacional Autónoma de México. Av. Universidad s/n, Col. Chamilpa, 62210. Cuernavaca, Morelos, México. ^3^Bioprocess Research Group, Department of Chemical Engineering, Universidad de Antioquia, Calle 70 No. 52-21, Medellín, Colombia.

### Supplementary results

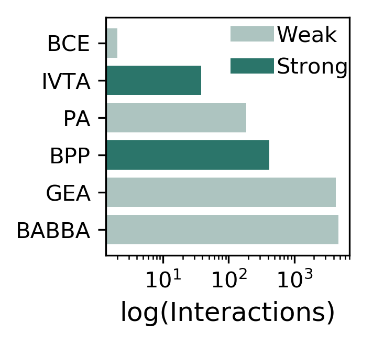

###### Supplementary Figure 1

The number of interactions supported by each type of experimental evidence. BCE, binding of cellular extracts; IVTA, in-vitro transcription assay; PA, proteomic analysis; BPP, binding of purified proteins; GEA, gene expression analysis; BABBA, binding affinity by bead-based assays.

The most studied TFs were the ones encoded by *phoP* (SCO4230) ^1-9^, *glnR* (SCO4159) ^10-16^, and the sigma factor encoded by *sigR* (SCO5216) ^17-23^. The interactions more studied are among the TF encoded by *phoP* (SCO4230) and its TGs *pstS* (SCO4142), *phoU* (SCO4228), *phoR* (SCO4229), and itself ^1,2,4,9^; and the sigma factor encoded by *sigR* (SCO5216) and its TGs *trxC* (SCO0885) ^19,20,22,23^, *trxB* (SCO3890), and itself ^18,20,22,23^ (Supplementary Table 3).

### The functional architecture of the *S. coelicolor* GRN

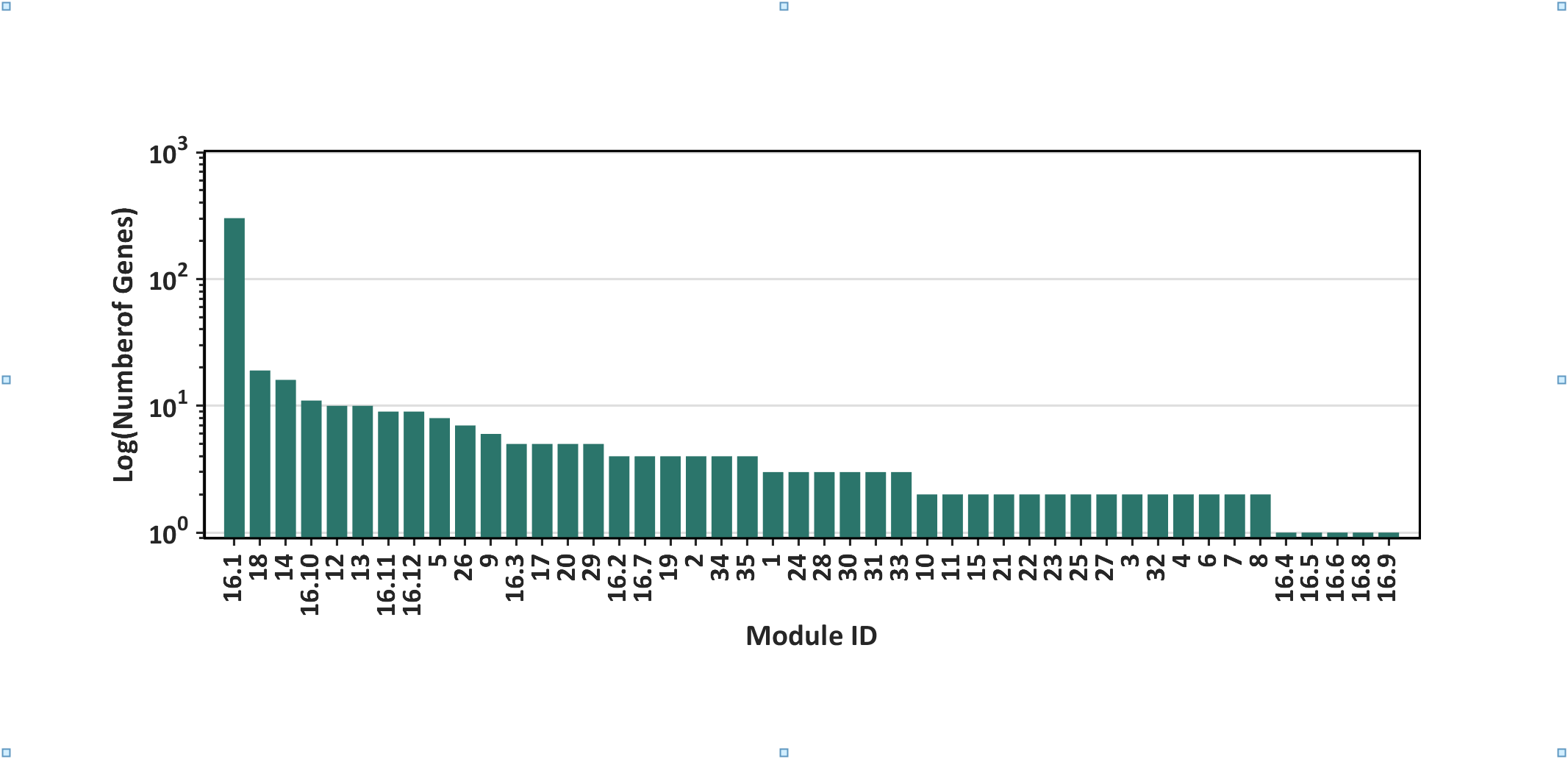

**a**

**b**

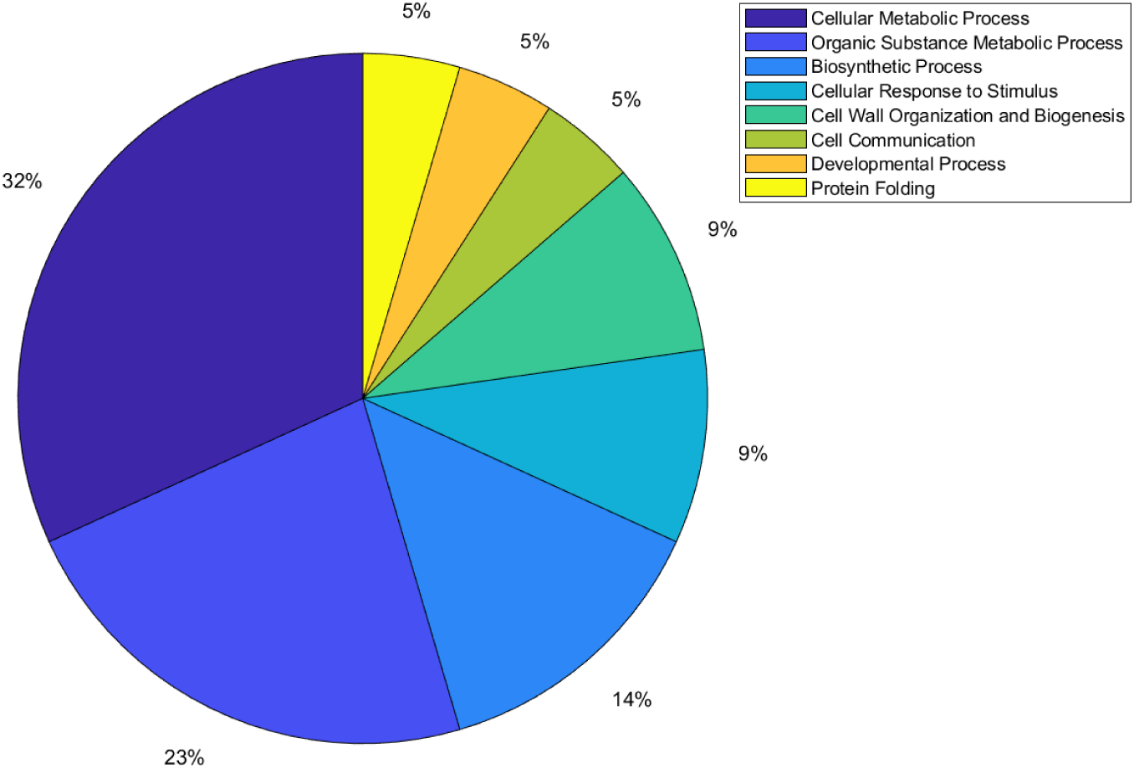

###### Supplementary Figure 2

**a)** Number of genes per module of the meta-curated network *Curated_RTB-FL-DBSCR.* **b)** Distribution of modules per function for the meta-curated network *Curated_FL-DBSCR-RTB.*

#### Global regulators

The nine GR previously reported in *Martín et. Al.*^24^ are *argR* (SCO1576), *absA2* (SCO3226), *phoP* (SCO4230), *afsS* (SCO4425), *abrC3* (SCO4596), *dasR* (SCO5231), *absC* (SCO5405), *ndgR* (SCO5552), and *scbR* (SCO6265). *phoP* is the gene with the highest out-connectivity in the meta-curated network *Curated_FL-DBSCR-RTB*. PhoP is a response regulator from the TCS PhoR–PhoP. It has been experimentally identified to act as GR in vivo controlling phosphate scavenging systems and cell wall/extracellular polymer biosynthesis ^9^. The other 10 genes that were classified as GRs by *Martín et. al,* were not identified as such by the NDA. The reason for these false negatives is the criteria used by the author to their classification since it is done by their capability to regulate genes from multiple pathways (wide domain regulators) or the regulation of hundreds of genes. Therefore, in an incomplete GRN, TFs controlling genes from multiple pathways but with a few TGs will not be identified as GRs by the NDA, where a high out-connectivity and low clustering coefficient of the gene are, by definition, features required to be classified as GR. Moreover, the classification by *Martín et. al* considers GRs for *Streptomyces* in general, not only for *S. coelicolor*. Therefore, genes labeled as false negatives could be acting as local regulators in *S. coelicolor* while being GRs in another *Streptomyces*.

Following, we describe the GRs or pleiotropic regulators that were reported individually (Supplementary Table 4). The sigma factor SigR (SCO5216) was recently reported as GR controlling DNA repair, protein quality control, thiol homeostasis, sulfur metabolism, ribosome modulation, and DNA repair^25^. ScbR2 (SCO6286) has been identified to regulate morphological differentiation and stress response through a plethora of genes across the *S. coelicolor* genome, suggesting a global-level regulation^26^. The ECF sigma factor SCO4117 has been previously reported as a pleiotropic regulator that controls secondary metabolism and morphogenesis ^27^. The gene SCO5283 has just been described to encode the cognate response regulator of the TCS SCO5282/SCO5283, having a pleiotropic effect in glycolysis, gluconeogenesis, stress-signaling pathways, proteins secretion, and cell envelope metabolism^28^. Rok7B7 (SCO6008) has been found to control carbon catabolite repression, antibiotic biosynthesis, xylose utilization, and morphological development^29^. The TF encoded by SCO7173 has been reported as pleiotropic regulators of phosphate starvation response and actinorhodin biosynthesis^30^. The TF encoded by SCO5785 is a response regulator related to antibiotic synthesis, sporulation, and several ribosomal genes ^31^. *Aor1* (SCO2281) has been recently described as a global regulator, orphan RR containing REC and HTH domains, which act as a positive regulator of antibiotic production of ACT, RED, and CDA; and of the genes involved in morphological differentiation^32^. WblA (SCO3579) has been reported as a pleiotropic regulator of various antibiotic pathways, the formation of aerial hyphae, and response to oxidative stress^33^. HrdB (SCO5820) is known to be the housekeeping sigma factor of *S. coelicolor* and to be essential for its survival, thus it affects a great number of biological processes and genes^34^. The only gene predicted as GR that has not been reported as such is SCO3356, which codes for the ECF sigma factor SigE and has only been reported as a coordinator of the cell wall integrity system^35,36^. Nevertheless, the maintenance of the wall integrity carries diverse biological functions, which might imply a pleiotropic effect on the cell.

#### Intermodular genes

Intermodular genes integrate the regulatory response of different modules, which means they coordinate different biological processes in the cell. Among all the curated networks, the meta-curated network *Curated_RTB-FL-DBSCR* has the largest number of intermodular genes, 18 (0.33% of the 5386 network genes). Some of the genes found as intermodular are: *ssgB* (SCO1541) a homolog of the sporulation gene *ssgA,* and its product has been suggested as a key regulator of the process of growth cessation before sporulation-specific cell division, affecting cell sporulation along with actinorhodin production^37^. Five genes are predicted as intermodular genes. Two genes *glnA* (SCO2198) and *glnII* (SCO2210) encode glutamine synthase and the *amount* operon, *amtB-glnK-glnD* (SCO5583-85), encodes an ammonium transporter, a PII protein, and an adenylyltransferase. These five genes are involved in nitrogen metabolism and are directed regulated by GlnR (SCO4159), the major regulator of nitrogen metabolism^6,38^. Moreover, they are also regulated by PhoP (SCO4230) the principal regulator of phosphate metabolism^6,38^. Moreover, *glnII* is also involved in the onset of mycelial differentiation, which might suggest a role in the regulation of secondary metabolism^39^. Additionally, an intrinsic role of glutamine synthetase in secondary metabolism has been also suggested in *S.* *lividans*^40^. This suggests that both genes might have in role in the coordination of nitrogen and phosphate metabolism, along with the secondary metabolism. Regarding another gene cluster *agl3EFG* (SCO7165-67) classified as intermodular, *Hillerich and Westpheling* suggested it have the function of carbohydrate transport^41^. This cluster appears to be regulated by GlnR (SCO4159) and Agl3R (SCO7168), a GntR family transcriptional regulator. While GlnR is known to regulate the genes involved in nitrogen metabolism^10^, many GntR family regulators have been proved to have a role of repression in carbon metabolism^41^. This might suggest that this cluster has a role in coordinating nitrogen and carbon metabolism, as it has been shown for other processes such as nitrogen and phosphate metabolism^42^. Other intermodular genes are the *actII-orf2* and *actII-orf3* (SCO5083-84), which are part of the BCG of ACT antibiotic. Both genes are regulated by *actII-orf4* (SCO5085) — the SARP of ACT —, AfsS (SCO4425) and SCO7173. -The two latter TFs are involved in phosphate metabolism and SCO7173 also affects the biosynthesis of ACT, which might be achieving through these intermodular genes^30^.

### Network inference

#### Quantitative assessment and cross-validation

**a	b**
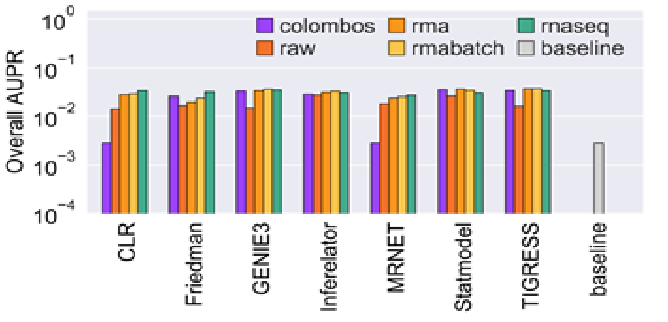

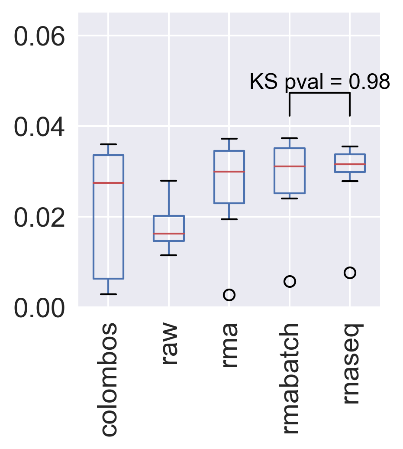

###### Supplementary Figure 3

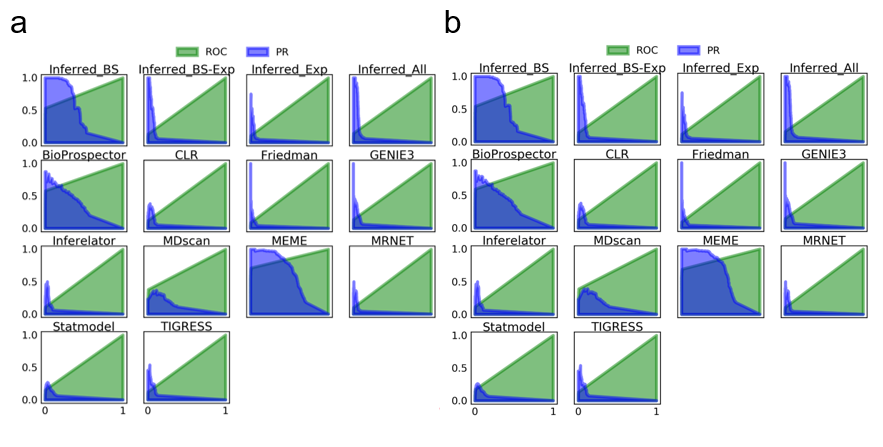
a) AUPR for the inferences from transcriptomic data. b) Distribution of the AUPR scores obtained with different datasets. KS = Kolmogorov-Smirnov between the distributions of the AUPR scores for batch-corrected RMA and RNA-seq datasets used as input for the GRNs prediction tools.

###### **Supplementary Figure 4**

ROC and PR curves for each of the inference methods and the community networks using the curated networks as GS.

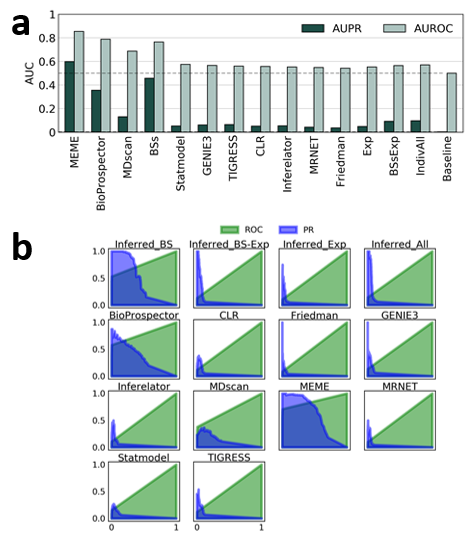

###### Supplementary Figure 5

a) AUCs for the inference methods and the community networks using the curated networks plus the statistically validated interactions as GS. b)ROC y PR curves for each of the inference methods and the community networks using the curated networks plus the statistically validated interactions as GS.

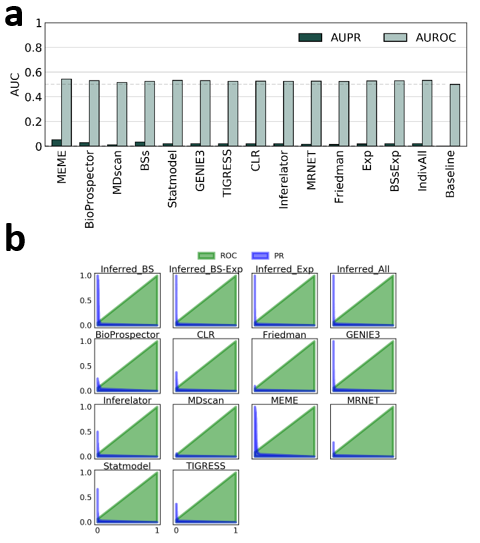

###### Supplementary Figure 6

a) AUCs for the inference methods and the community networks using every curated interaction as the GS (Curated_FL-DBSCR15-RTB13). Note that the tools based on binding sites prediction are the most affected ones wrt. Supplementary Figure 5. This is because the predictions based on binding sites were only for the experimentally supported DNA-binding proteins. Namely, the TFs in Curated_FL(S)-DBSCR(S). Curated_FL(S)-DBSCR(S) has only 72 TFs, while Curated_FL-DBSCR15-RTB13 has 137. This results in a poor recall by the predictions based on binding sites (b), as interactions for 65 TFs are not predictable. Even though the precision increased for most of the tools based on gene expression data, they also showed a poor recall.

##### Topological assessment

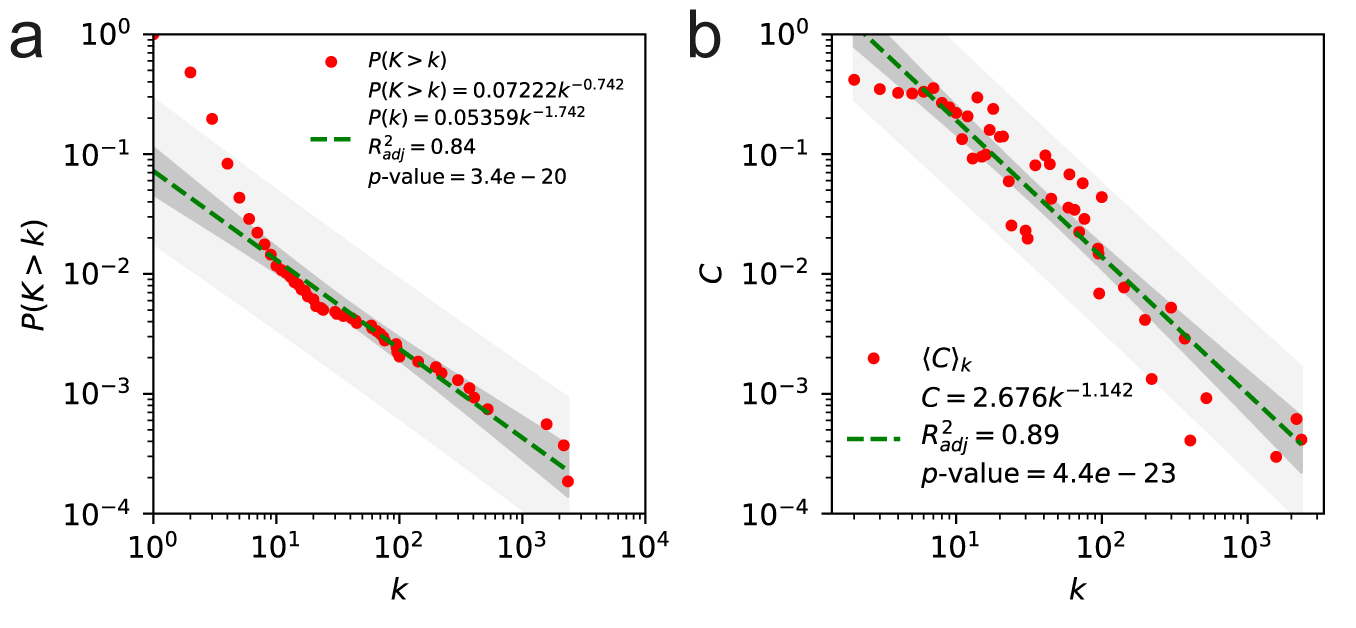

###### Supplementary Figure 7

a) Cumulative distribution of node connectivity and b) clustering coefficient distribution for the meta-curated network *Curated_FL-DBSCR-RTB.*

Some of the most relevant structural properties of the curated networks are described following. The curated networks have low network density (<1%), as have been previously reported for other bacterial regulatory networks **^43^**. Their average path lengths are smaller than the logarithm of the network size, $\ln(ln(N))$, which indicates that they are ultra-small world networks, as scale-free networks **^44^**. From the robust linear regression over the log-log data, the degree distributions ($P(k)$) follow a power-law, suggesting that they are scale-free (Supplementary Figure 7). **Regarding** the clustering coefficient distribution ($C(k)$), the networks seemed to be hierarchical modular. Both characteristics were previously observed in diverse bacterial networks **^45^**. We computed the Kolmogorov-Smirnov distance between the networks’ degree distribution and several probability functions, and a maximum likelihood estimation for their α to corroborate that the curated networks are scale-free. Except for Curated_DBSCR(S) that has a similar distance between its degree distribution to a power law and a log-normal, the degree distributions of all the curated networks have the smallest distance to a power law distribution (Supplementary Table 7a). For the α estimation values between two and three for the curated networks (Supplementary Table 7b), we can corroborate that the curated networks are scale-free as other bacterial GRNs **^43^**

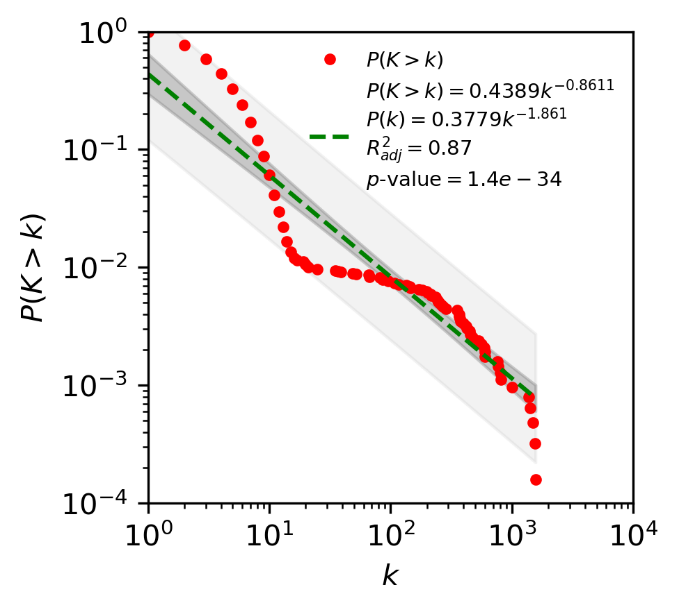

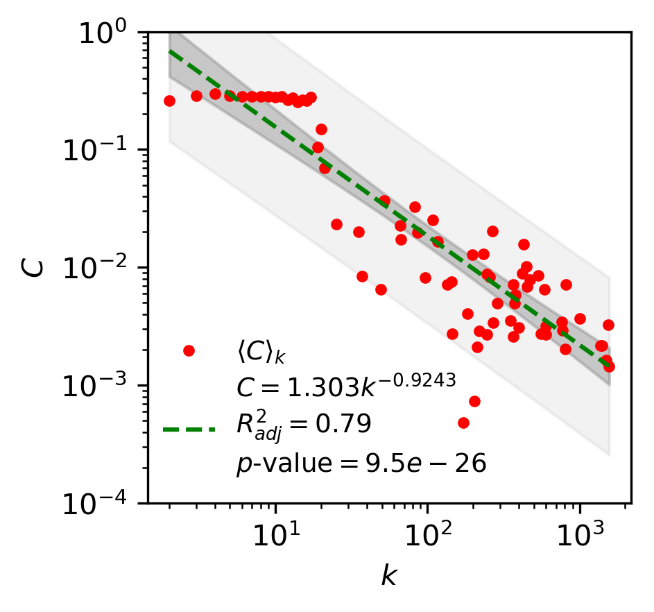

###### Supplementary Figure 8

Cumulative distribution of node connectivity and distribution of node clustering coefficient of community network *Inferred_BS.*

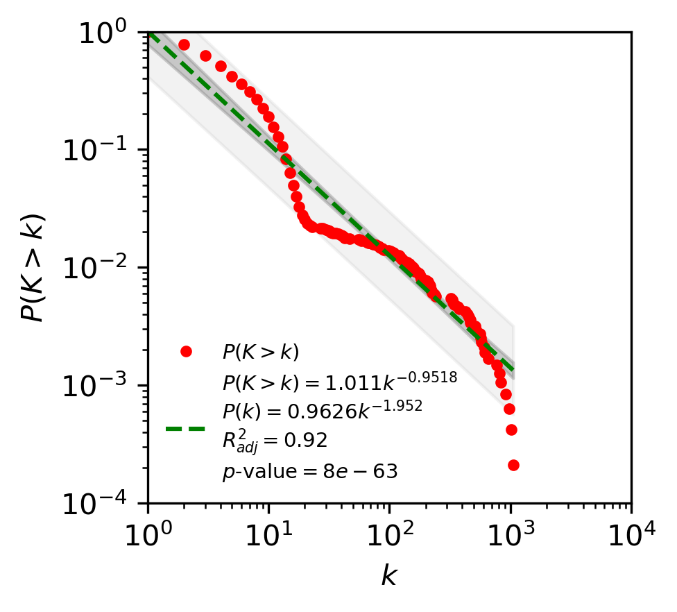

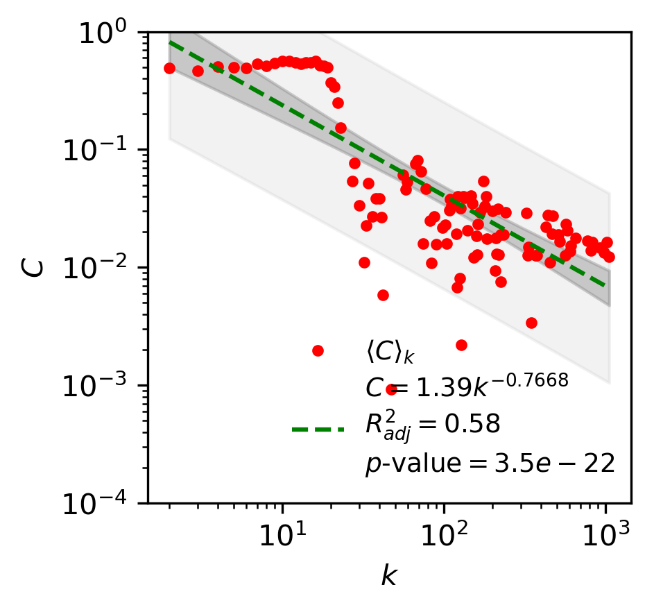

###### Supplementary Figure 9

Cumulative distribution of node connectivity and distribution of node clustering coefficient of community network *Inferred_Exp.*

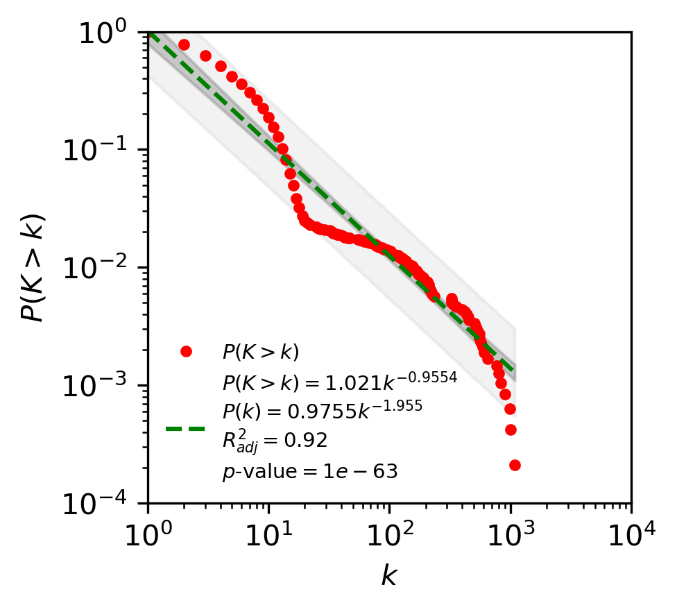

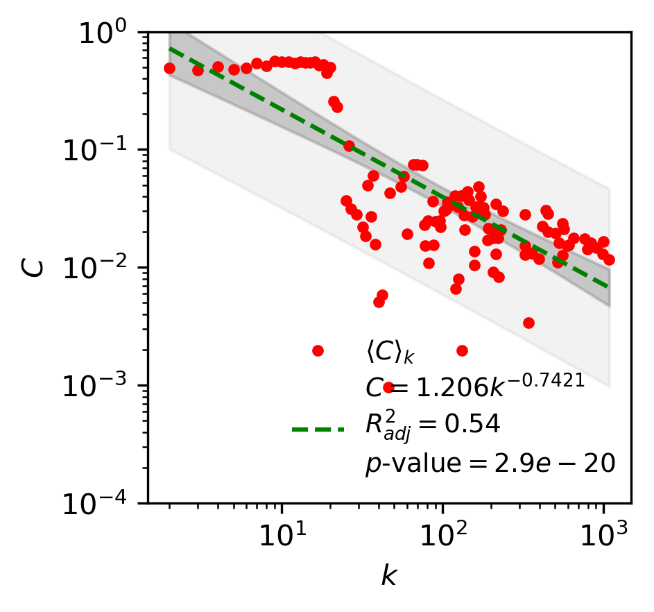

###### Supplementary Figure 10

Cumulative distribution of node connectivity and distribution of node clustering coefficient of community network *Inferred_BS-Exp.*

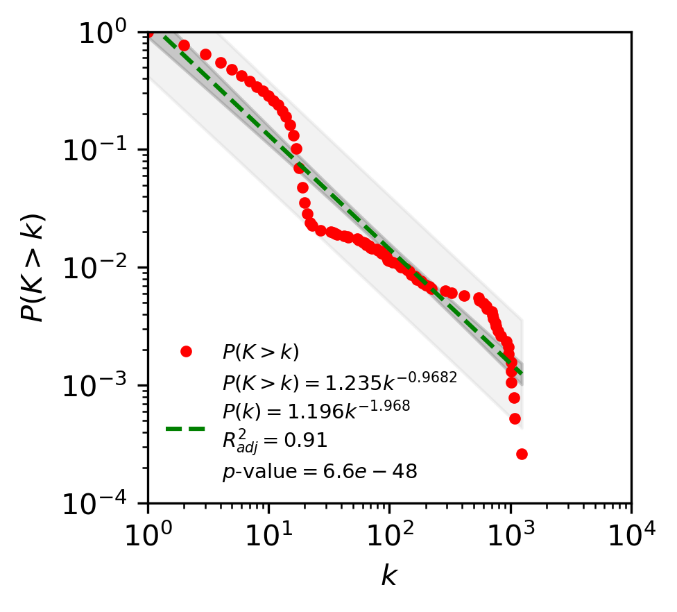

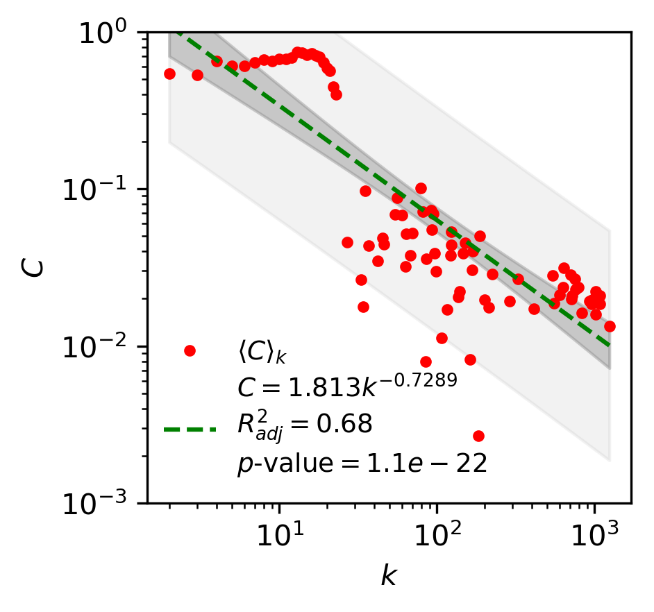

###### Supplementary Figure 11

Cumulative distribution of node connectivity and distribution of node clustering coefficient of community network *Inferred_All.*

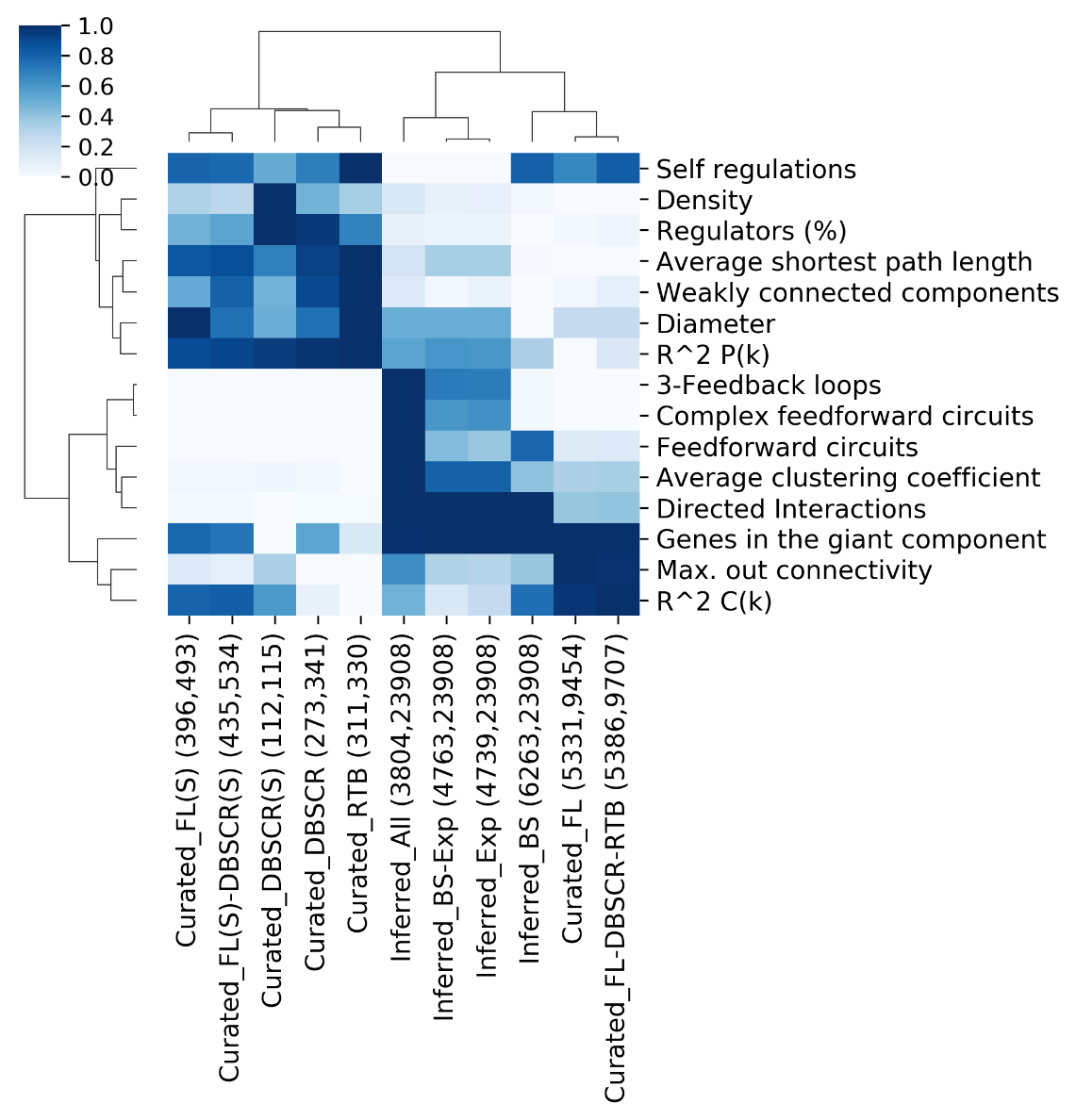

###### Supplementary Figure 12

Comparative of network properties. Min-max normalization was applied for each property across networks. Aside from each network is their cognate number of nodes and interactions are displayed.

##### Assessment of the global regulators' inference

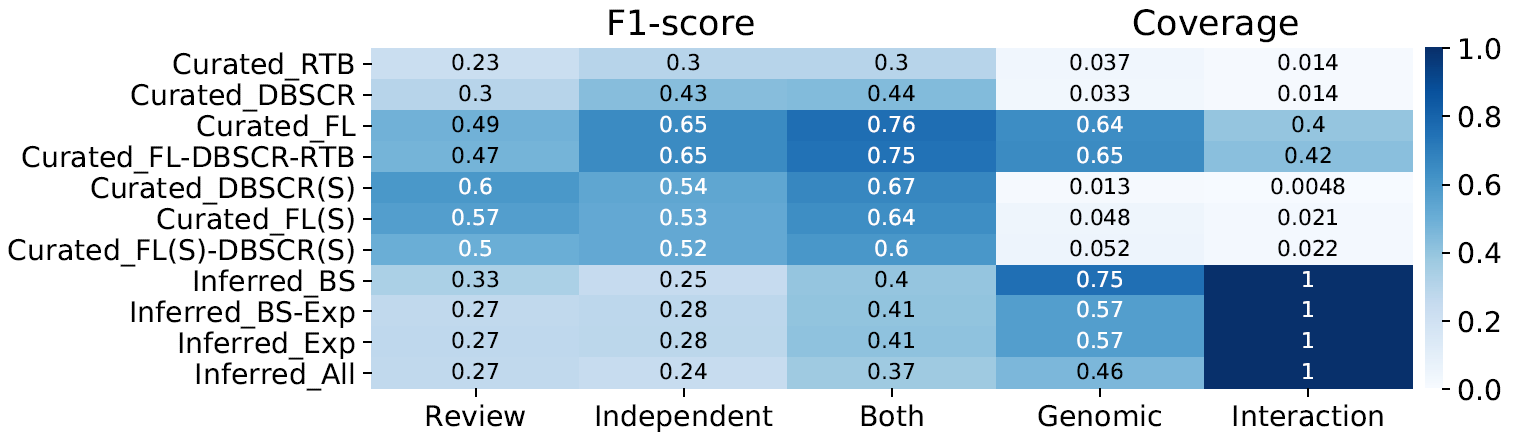

###### Supplementary Figure 13

F1-score for global regulators predicted by NDA for each of the curated and inferred networks. Scores ≥ 0.5 are represented in white numbers.

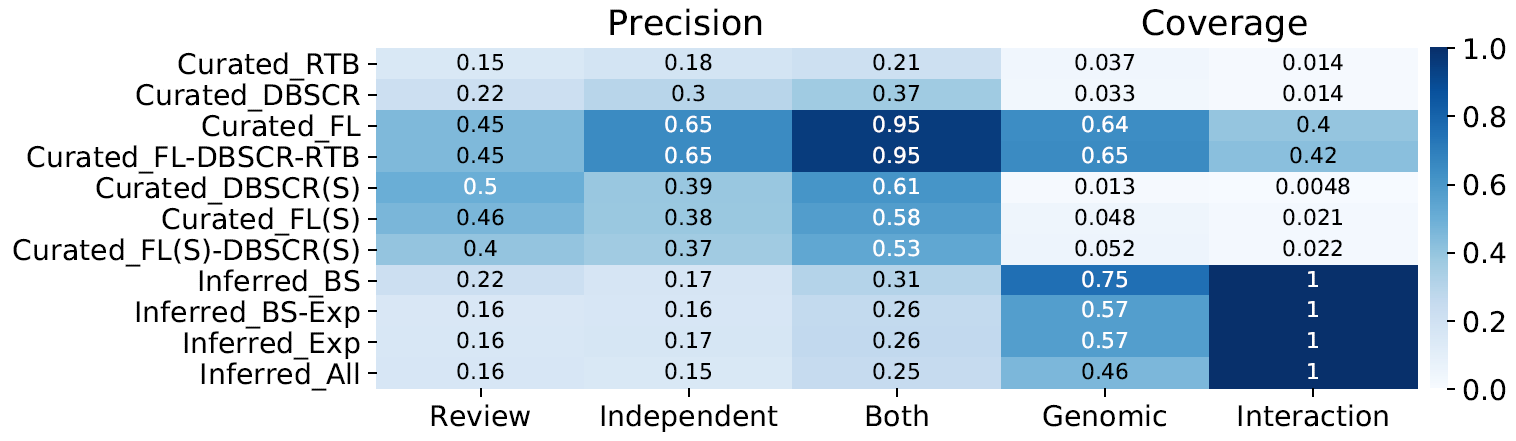

###### Supplementary Figure 14

Precision for global regulators predicted by NDA for each of the curated and inferred networks. Scores ≥ 0.5 are represented in white numbers.

##### Putative transcription factors for *actII-orf4*

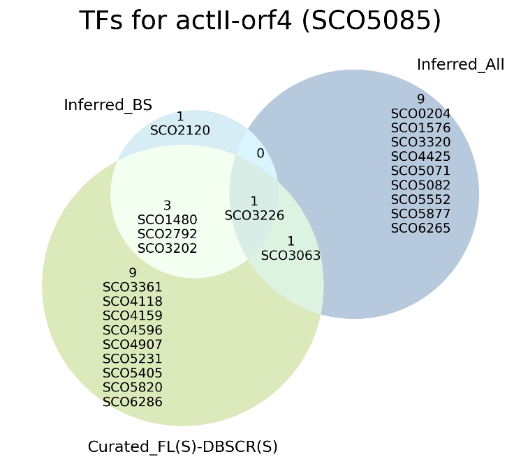

###### Supplementary Figure 15. Potential transcription factors for *actII-orf4* suggested by *Inferred_BS* and *Inferred_All*, and the overlap of their predictions with the strong meta-curation.

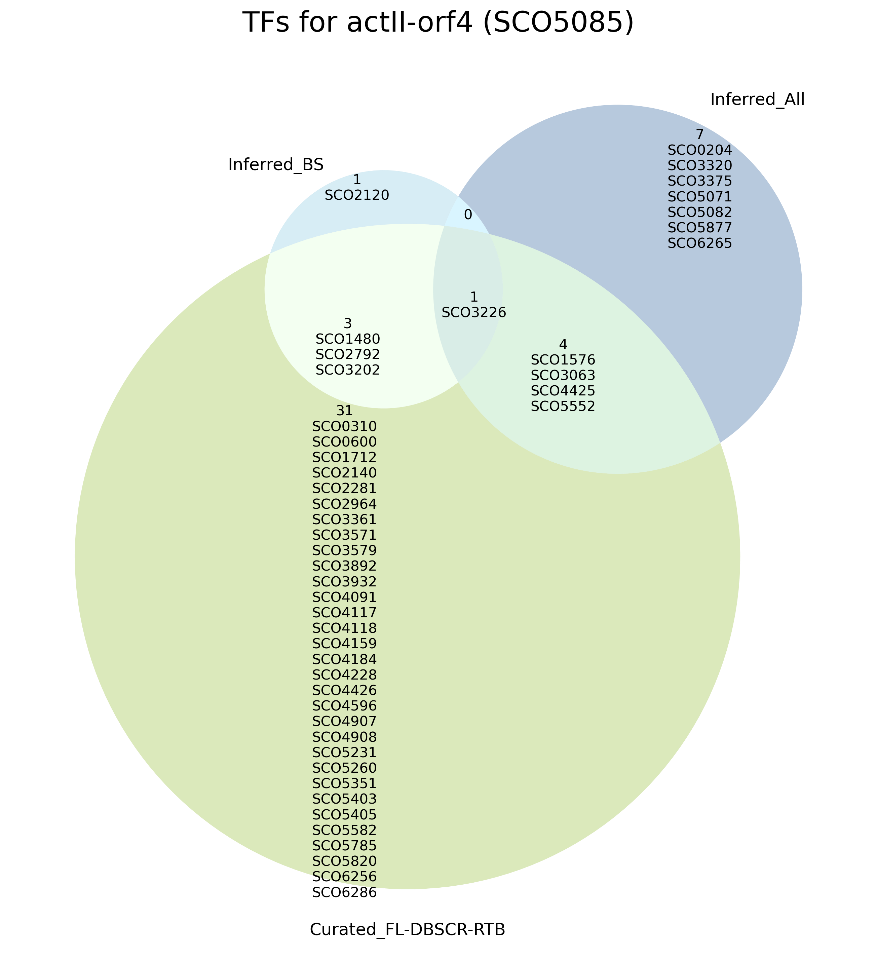

###### Supplementary Figure 16. Potential transcription factors for *actII-orf4* suggested by *Inferred_BS* and *Inferred_All*, and the overlap of their predictions with the meta-curation including all types of experimental evidence.

- Besides SCO3226 and SCO3063, 3 more TFs have been already reported as regulators of SCO5085 and are included in the network *Curated_FL-DBSCR-RTB*. Namely, SCO1576, SCO4425, and SCO5552.
- SCO0204 (*osdR*) regulates genes involved in the control of stress and development^46^. The authors characterized the OsdR regulon using predictions based on regulatory binding sites and verify them with EMSA. However, SCO5085 was not a putative target in their prediction. It was not part of our prediction based on binding sites (*Inferred_BS*) neither, only of predictions based on gene expression.
- SCO3320. Previous work suggested the regulation of SCO5085 by SCO3320 through a LexA-like motif in *E. coli*^47^. However, no experimental evidence has been performed to confirm such regulation.
- SCO3375 codes a putative Lsr2-like protein regulated by PhoP and regulates the transcription of SCO1541, a gene involved in sporulation-specific cell division.
- SCO5071 codes a putative hydroxylacyl-CoA dehydrogenase and has been found its expression to be regulated by SCO5351^48^, a regulator of SCO5085. Noteworthy, SCO5071 has been reported as a TG for SCO5085 itself ^49^. SCO5071 is included as a regulator of several genes in *Curated_RTB*, however, it might be an indirect effect such as the feedback expected to happen with SCO5085.
- SCO5082 codes ActII-orf1, a repressor of *actII-orf2* and *actII-orf3*^50^. The genes *actII-orf2* and *actII-orf3* are divergently transcribed, therefore, it is likely for *actII-orf4* to also be regulated by ActII-orf1.
- SCO5877 codes RedD, a regulator of the RED biosynthesis^51^. Suggesting cross-regulation between gene clusters encoding biosynthesis of individual antibiotics^52^.
- SCO6265 codes ScbR, a protein involved in γ-butyrolactone (SCB1) synthesis that has been previously suggested to regulate Act and Red synthesis through *orfB* (SCO6268)^53^.

### Supplementary methods

#### Details collected in the curation of the regulatory interactions

1. TF
2. TF locus tag
3. TF description
4. TG
5. TG locus tag
6. Experiment
7. evidence (methodology applied for the identification of the interaction)
8. regulatory function (repression, activation, unknown)
9. evidence classification, interactions were classified according to the weight of the experimental evidence supporting them as “strong” (evidence of direct TF binding) or “weak” (no evidence of direct TF binding), following the RegulonDB scheme ^54,55^ (Supplementary Figure 1)
10. reference PubMed ID and DOI of the paper curated; xi) year of publication; xii) notes.
11. The same information is presented in Supplementary Table 1.

#### Transcriptomic Data

First, we used the microarray data consisting of 371 samples available from the COLOMBOS Database^56^. Second, we obtain microarray and RNA-Seq data from NCBI GEO^57^. As there is not a consensus method for microarray data integration from different platforms, we decide to take the platform with the largest dataset (137 samples), which was an Affymetrix platform. This data was taken raw, then Robust Multi-chip Averaging (RMA) normalization was performed^58^, and finally, it was batch-effect corrected^59^. For the case of RNA-Seq, the data is available with different types of normalization for each series. Therefore, we decided to take the largest dataset as well (54 samples). The inference from expression data was performed over the 5 datasets available: i) COLOMBOS (colombos), ii) RNA-Seq (rnaseq), iii) Affymetrix raw (raw), iv) Affymetrix with RMA normalization (rma), and v) Affimetrix with batch-effect correction (rmabatch). To provide insights on the quality of the predictions, the inferred GRNs were assessed using the network Curated_FL(cS)-DBSCR(S) as GS, and the AUPR were computed for each one of the networks. We assessed the inferred GRN based mostly on the AUPR since it is more informative for imbalanced datasets^60^ as it is the case of GRNs inference^61^ (Supplementary Figure 3). From the evaluation, despite a large amount of data, the prediction with colombos performs poorly regarding the other data sets. From the Affymetrix data, the rmabatch set performed the best with a very similar result to the rnaseq dataset. However, the latter comes from one unique study, while the Affymetrix data comes from different studies with more diverse data. Then, we selected the Affymetrix rmabatch as our dataset for the final inference from transcriptomic data. This, since we believe a more diverse data will allow us to identify a higher quantity of regulatory interactions.

#### Friedman

We proposed a modification of the method ANOVA for GRN inference. In this method, the likelihood of an interaction between a transcription factor (TF) and a target gene (TG) is given by a non-linear correlation coefficient, which is derived from an analysis of variance (two-way ANOVA). The correlation coefficient measures association as the fraction of the total variance that is explained by the differential expression across experimental conditions ^62^. ANOVA has some requirements for its proper application, one of them is a normal distribution. This might not be accurate for the distributions of gene expression. Thus, we proposed a variation, deriving the non-linear correlation coefficient from a Friedman test. This is the non–parametric alternative of ANOVA since it does not make assumptions of normality ^63^. A Matlab implementation for the original method (Anova) and the modification (Friedman), along with their documentation can be found at <https://github.com/andreazorro/Anova>. The algorithm takes as input a matrix of expression data and a list of TFs, and the output is the list of TF - TG correlation coefficients.

#### Statmodel

We proposed a modification for GRN inference of the method Statmodel proposed by Hernandez^64^. This method is an alternative tool for statistical modeling and analysis of experiments than ANOVA. Since this methodology allows us to determine the influence of independent factors (TFs expression) over a response variable (TG expression), we found it as a proper tool for GRN inference. For this purpose, we proposed a modification of the methodology initially presented and a suggested score for the interaction reliability. Statmodel presents some advantages concerning ANOVA, such as no assumption of the data distributions and the minimization of the variance of the residual error probability model, reducing the chances of over-fitting. Moreover, this methodology reduces the spurious effects minimizing the number of predictor variables, which is mainly what we look for in GRN inference.

To obtain a parsimonious model, we should reduce the number of parameters of the model. This can be done through a hypothesis test to find the coefficients that are significantly different from zero. To perform the hypothesis test, first, we find the probability distribution that better adjusts to the response variable. Then, we evaluate if the data for the smallest subgroup is adjusted to this probability distribution (See Hernandez^64^ for further details). To evaluate this, we applied a Chi-square test in Matlab. The coefficient is considered to be significantly different from zero If the alternative hypothesis cannot be rejected. This means that the null hypothesis Is rejected by the Chi-square test.

In this case, we are evaluating the influence of a TF's expression on a gene’s expression. We proposed a score for the reliability of the interaction as the $-\log\left( \text{p-value} \right)$ of Chi-square test. Its Matlab implementation along with its documentation can be found at <https://github.com/andreazorro/Statmodel>. The algorithm takes as input a matrix of expression data and a list of TFs, and the output is the list of TF-TG reliability scores.

#### The Natural Decomposition Approach

First, the NDA identifies the global regulators (GRs) as genes with higher out connectivity than the κ value, the equilibrium point from the $C(k)$ distribution where the variation of the clustering coefficient is equal to the variation of the connectivity but with the opposite sign *dC*(*k*)/*dk* = -1. Then, the genes from the basal machinery are revealed as those regulated only by GRs. After the removal of the GRs and the genes from the basal machinery, as well as their interactions, the structural genes are removed, leaving isolated subnetworks (modules). Then, the structural genes are re-integrated as intermodular genes if their regulators belong to different modules, otherwise as modular genes belonging to the same module as their regulators.
